# Mutation and selection induce correlations between selection coefficients and mutation rates

**DOI:** 10.1101/2023.02.13.528299

**Authors:** Bryan L. Gitschlag, Alejandro V. Cano, Joshua L. Payne, David M. McCandlish, Arlin Stoltzfus

**Author notes:** These authors contributed equally (Bryan L. Gitschlag, Alejandro V. Cano).

## Abstract

The joint distribution of selection coefficients and mutation rates is a key determinant of the genetic architecture of molecular adaptation. Three different distributions are of immediate interest: (1) the *nominal* distribution of possible changes, prior to mutation or selection, (2) the *de novo* distribution of realized mutations, and (3) the *fixed* distribution of selectively established mutations. Here, we formally characterize the relationships between these joint distributions under the strong selection, weak mutation (SSWM) regime. The *de novo* distribution is enriched relative to the nominal distribution for the highest rate mutations, and the fixed distribution is further enriched for the most highly beneficial mutations. Whereas mutation rates and selection coefficients are often assumed to be uncorrelated, we show that even with no correlation in the nominal distribution, the resulting *de novo* and fixed distributions can have correlations with any combination of signs. Nonetheless, we suggest that natural systems with a finite number of beneficial mutations will frequently have the kind of nominal distribution that induces negative correlations in the fixed distribution. We apply our mathematical framework, along with population simulations, to explore joint distributions of selection coefficients and mutation rates from deep mutational scanning and cancer informatics. Finally, we consider the evolutionary implications of these joint distributions together with two additional joint distributions relevant to parallelism and the rate of adaptation.

## Introduction

Evolutionary change has long been understood as a process combining variation and selection. Within this duality, a range of views is possible. Darwin theorized that variation merely supplies abundant materials, with no dispositional influence: selection does all the important work of choosing, so that the laws of variation “bear no relation” to the outcomes built by selection (Darwin, 1868, Ch. 21). By contrast, a century later, when sequence comparisons revealed a process of change that reflects both mutation and selective filtering, selection was depicted as the editor, not the composer, of the genetic message (King and Jukes, 1969). The ability of mutation to shape patterns of molecular divergence was initially treated as an aspect of neutrality, but more recent work has shown that, when adaptation involves new mutations, differences in the rate that specific variants are introduced into the population can have a strong influence on the genetic basis of adaptation (Bailey et al., 2017; Cano et al., 2023, 2022; Couce et al., 2015; Gomez et al., 2020; Rokyta et al., 2005; Sackman et al., 2017; Schenk et al., 2022; Stoltzfus, 2021; Stoltzfus and McCandlish, 2017; Storz et al., 2019).

Because of the powerful and long recognized influence of selection on the changes involved in adaptation, the distribution of fitness effects (DFE) for mutationally possible changes has long been regarded as a fundamental parameter in evolutionary genetics (Eyre-Walker and Keightley, 2007; Kassen and Bataillon, 2006; Orr, 2003; Sanjuan et al., 2004). Yet, the role of mutation rates suggests the importance of, not merely a distribution of selection coefficients, but a combined distribution of mutation rates and fitness effects, i.e., the joint distribution of selection coefficients and mutation rates for a set of possible changes. This need is implicit, for instance, in treatments of the genetic architecture of adaptation that address genotypic redundancy (Láruson et al., 2020), distinguishing the case where several redundant mutations share the same selection coefficient from the case of a single mutation produced at the same total rate. While one might suppose that mutation rates and selection coefficients are inherently uncorrelated and thus the joint distribution is of little interest, here we show that this is not the case at all: even if mutation rates and selection coefficients are uncorrelated among possible beneficial mutations, the process of adaptation can induce correlations in the realized distribution of adaptive changes due to the heterogeneity of mutation rates and the enrichment by selection of mutations with large fitness effects. Furthermore, substantial correlations between mutation rates and selection coefficient among possible beneficial mutations can emerge merely due to having a finite mutational target size, particularly when only a small number of beneficial mutations are available.

Here we analyze the joint distribution of mutation rates and selection coefficients among beneficial mutations, and provide a formal treatment of the correlation between mutation rates and selection coefficients induced by heterogeneity in mutation rate and the probability of fixation. Fig. 1 gives a schematic illustration of a case where there are 40 possible beneficial mutations, with mutation rates and selection coefficients that are uncorrelated (Fig. 1, left). Whereas this distribution over possible beneficial mutations, which we call the *nominal* distribution, weights all mutations equally, mutations with higher mutation rates enter the population more frequently, producing a second joint distribution for *de novo* mutations that is reweighted linearly according to mutation rate, thus shifting the density to the right, toward mutations with higher rates (Fig. 1, center). We see in this case that mutation rate and selection coefficient are negatively correlated among *de novo* mutations even though they were uncorrelated among all possible beneficial mutations.

**Fig. 1.**
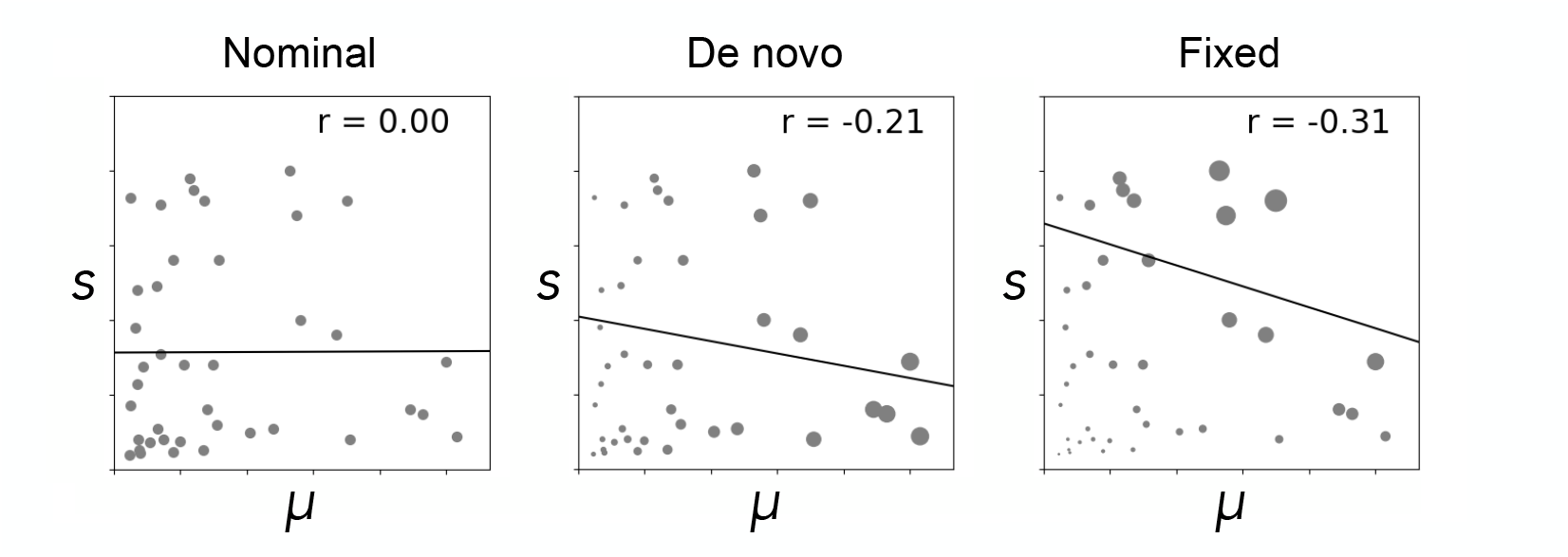
Mutation and selection reweight the joint distribution of mutation rates and selection coefficients. Three joint distributions of mutation rate, *μ*, and selection coefficient, B, (both in arbitrary units) are shown. The nominal distribution shows the set of possible beneficial mutations. The production of variation by mutation yields a *de novo* distribution weighted by mutation rate, shifting the density to the right. The fixed distribution reflects both this rightward shift and an upward shift due to selection. Such shifts can result in substantial correlations in the *de novo* and fixed distributions even if there is no correlation in the nominal distribution.

Finally, the action of natural selection favors mutations with higher selection coefficients, transforming the *de novo* distribution into the expected joint distribution of selection coefficients and mutation rates among fixed mutations (Fig. 1, right). In this case we assume that the probability of fixations is given by 2B (Haldane, 1927), as would be the case in the strong selection weak mutation (SSWM) regime, so that this distribution is derived from the *de novo* distribution by linearly re-weighting mutations according to their selection coefficient. We see that besides shifting the distribution of fixed mutations towards mutations with higher selection coefficients, this re-weighting also produces an even stronger negative correlation between selection coefficients and mutation rates among fixed mutations, despite the fact that mutation rates and selection coefficients were initially uncorrelated.

We proceed as follows with a more general treatment of this problem. First, we define the distributions of interest, and explain the relations between them, which can be captured using the concept (from probability theory) of a size-biased distribution (Arratia et al., 2019). We then describe a simple model of mutagenesis and adaptation in which the possible mutation rates and selection coefficients each take on just two or three discrete values. Using this model, which is easily visualized, we show that the correlation between mutation rates and selection coefficients can exhibit any pattern of signs across the three distributions. We then derive a more general theory of joint size-biasing via mutation and selection, and use this model to explain why drawing mutation rates and selection coefficients from independent exponential distributions tends to result in a negative correlation between selection coefficient and mutation rate among fixed mutations.

To illustrate how these ideas may be applied to actual cases, we then consider experimental and clinical data, beginning with high-throughput data on mutation and fitness for Dengue virus variants from Dolan et al. (2021). For beneficial variants, we show that the resulting fixed distribution shows a negative correlation between mutation rates and selection coefficients, with a magnitude that depends on the mutation supply (*N μ*). We also consider empirical data on the joint distribution of mutation rates and selection coefficients based on a deep mutational scanning study of the TP53 gene (encoding tumor protein p53), which allows for the construction of a nominal distribution of potential changes to TP53. We then use clinical prevalence data for tumor-associated mutations to represent the fixed distribution, i.e., a distribution reflecting both mutation and selection, revealing relatively modest effects on the fixed distribution. A stronger effect is seen when tumor-associated mutations are separated into those arising by single-nucleotide or multi-nucleotide mutations: the multi-nucleotide mutations that are observed clinically, in spite of lower mutation rates, tend to have larger selection coefficients.

We conclude with a prescription for future work. For mutation-limited evolution, the theory of mutation-fitness associations describes how a potential distribution of possibilities is transformed into an actual one. The data necessary to evaluate this theory empirically is increasingly available, as technological advances continue to facilitate the large-scale measurement of mutation rates and selection coefficients. Further work is needed to understand what kinds of transformations are most likely to occur in nature, and what observed transformations can tell us about underlying adaptive processes.

## Analytical results

Here we consider the joint distribution of mutation rates and selection coefficients for beneficial changes from three different perspectives. First, we can ask about the joint distribution of selection coefficients and mutation rates when we pick uniformly from some set of possible mutations. For example, if we have a DNA sequence of length *ℓ*, then there are 3*ℓ* alternative sequences that differ by a single nucleotide, out of which some number *n* are beneficial. Any one of these *n* beneficial changes occurs by mutation at some definite rate and has some definite selective advantage. Accordingly, we could ask about the expected selection coefficient *E* (*s*), for instance, or the correlation between mutation rate and selection coefficient, when we draw from these *n* possibilities uniformly at random. We call this the *nominal* distribution and denote the expected selection coefficient or mutation rate of a draw from this distribution as *E*_nom_(*s*) or *E*_nom_(*μ*), respectively. The nominal distribution corresponds to the results of a typical deep mutational scanning study that measures the effects of all single-nucleotide variants (or in other designs, all amino acid variants), or when we scan a model of context-dependent mutation rates across the genome and calculate the mutation rate for each possible mutation. This is also the distribution relevant for determining the overall mutation rate, since the total beneficial mutation rate is the sum of the mutation rates to each possible beneficial alternative.

Second, in addition to the joint distribution of selection coefficients and mutation rates among mutational possibilities, we can consider the distribution of spontaneously occurring mutations. Here, instead of picking (for instance) a random position and a random nucleotide, as in the nominal distribution, we ask about the distribution of the next beneficial mutation to occur. This distribution differs from the nominal distribution in that mutations with higher rates are more likely to be the next mutation to occur. This is also the distribution that we observe in mutation accumulation experiments, and the distribution relevant to determining the average selection coefficient among new mutations. We call this the distribution of *de novo* mutations and write the expected selection coefficient or mutation rate of a new mutation as *E*_de novo_(*s*) or *E*_de novo_(*μ*).

Third, we can consider mutations that become fixed in evolution, that is, those that actually contribute to adaptation. This is equivalent to asking about the selection coefficient and mutation rate of the next variant that is going to fix in the population. We call this distribution the *fixed* distribution and write the expected selection coefficient or mutation rate of the next mutation to fix as *E*_fixed_(*s*) or *E*_fixed_(*μ*).

To understand the necessary relationships between these distributions, it is helpful to introduce the concept of a size-biased distribution (Arratia et al., 2019). In particular, given a non-negative random variable *X*, we can define the size-biased distribution as the random variable *X*^*^, where *P*(*X*^*^ = *x*) is proportional to *xP*(*X* = *x*) for any non-negative value *x*. Thus, the size-biased distribution is a re-weighted version of the original probability distribution. More specifically, it is the probability distribution obtained when the new weights on each outcome are given by the value of that outcome. This re-weighting favors larger outcomes and results in a systematic change to the moments of the distribution.

More precisely, the *k*^th^ raw moment of the size-biased distribution is determined by the (*k* + 1)^th^ moment of the original distribution by *E* ((*X*^*^)^*k*^) = *E* (*X*^*k*+1^)/*E* (*X*). Size-biasing is well known to result in certain counter-intuitive phenomena (Ross, 2014, Section 7.7). For example, if the number of children per family is given by the random variable *X* then the number of children in a random child’s family is given by *X*^*^, so that the average number of siblings of a random child is greater than would be expected based on the size of an average family. Another classical example is that, if the time interval between buses is distributed as *X*, the expected time spent waiting for a bus, for a person who arrives at the bus stop at a random time, is *E* (*X*^*^)/2 instead of the shorter time *E* (*X*)/2, because a person is more likely to arrive in a longer interval between buses than a smaller one.

The concept of size-biasing is helpful in thinking about the relationships between the nominal, *de novo*, and fixed distributions because these distributions are related to each other by size-biasing according to either the mutation rate or the selection coefficient. In particular, the *de novo* distribution is obtained by size-biasing the nominal distribution with respect to the mutation rate. In symbols, we write this as:

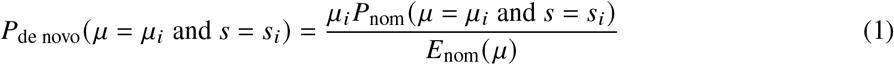

where *s*_*i*_ and *μ*_*i*_ are the selection coefficient and mutation rate for mutation *i*, respectively.

This size-biasing means that the average mutation rate of mutations observed in the *de novo* distribution will be at least as large as the average mutation rate with respect to the nominal distribution. In fact, using Var to denote the variance, the increase in the expected mutation rate is given by

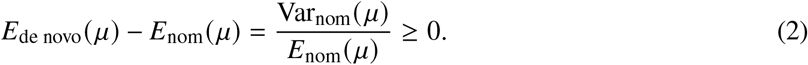

Importantly, when a non-zero correlation exists between mutation and selection in the nominal distribution (a situation that, as argued below, often occurs by chance when the number of possible beneficial mutations is small or moderate), size-biasing the nominal distribution with respect to mutation rate will also affect the mean selection coefficient. In particular, using Cov for the covariance,

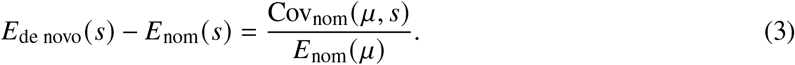

Equations 2 and 3 are an immediate consequence of more general results on the effects of size-biasing that are derived in the Appendix as Equations A2 and A5, respectively. Indeed, the form of these equations will likely be familiar to many readers as they are directly analogous to Fisher’s fundamental theorem and Robertson’s secondary theorem, respectively (see Queller 2017). This is because the fitness distribution after selection is the size-biased transformation of the fitness distribution before selection, and the response to selection corresponds to size-biasing the distribution of another phenotype (a second random variable) according to its fitness. Thus, these classical results for changes due to selection in a single generation can likewise be derived from Equations A2 and A5, respectively, and provide an easy mnemonic for the results of size-biasing more generally.

Whereas the *de novo* distribution is obtained from the nominal distribution by size-biasing with respect to mutation rate, the fixed distribution is obtained from the *de novo* distribution by re-weighting with respect to the probability of fixation. Although in general the probability of fixation *π* is a function of the entire *de novo* distribution as well as other population-genetic parameters such as the population size (e.g., see Gerrish and Lenski, 1998; McCandlish and Stoltzfus, 2014; Neher, 2013), here we assume conditions, explained as follows, under which this re-weighting can be treated as a case of size-biasing proportional to *s*. As the mutation supply *Nμ* becomes small, new mutations enter the population (i.e., originate) one at a time and each allele can be assigned an individual probability of fixation *π*_*s*_. This is called the origin-fixation regime or, when considering beneficial alleles with *s* ≫ 1/*N*_*e*_, the strong-selection, weak-mutation (SSWM) regime (Gillespie, 1994; McCandlish and Stoltzfus, 2014). For a Wright-Fisher population, the probability of fixation per Haldane (1927) is *π*_*s*_ ≈ 2*s* when *s* is small relative to 1, and a probability of fixation proportional to *s* holds for a wide class of models beyond the Wright-Fisher model (Patwa and Wahl, 2008), so long as *s* is small relative to 1 and large relative to 1/*N*_*e*_. Thus, our mathematical approach based on size-biasing proportional to *s* corresponds precisely to this set of mutation-limited conditions where the fixation of beneficial mutations is proportional to *s*, and so our mathematical results are valid for those conditions. In a more complex scenario in nature or in the population simulations below, the emergence of the fixed distribution may reflect non-proportionality due to clonal interference and saturation of the probability of fixation as it approaches 1 when *s* becomes large.

More formally, assuming that the probability of fixation is a linear function of the selection coefficient, we have

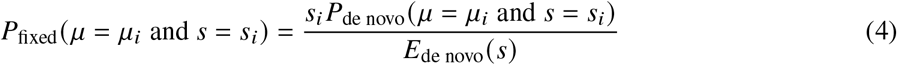

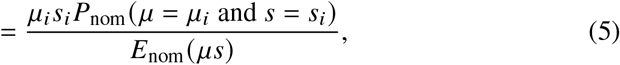

where the second line shows that we can also derive the fixed distribution from the *de novo* distribution by reweighting each class of mutations 8 according to the product of its mutation rates and selection coefficients *μ*_*i*_ *s*_*i*_. Similarly to the difference between the nominal and *de novo* distributions, moving from the *de novo* distribution to the fixed distribution will typically change the expected selection coefficient and expected mutation rate, with

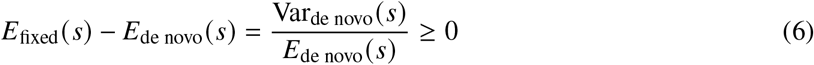

and

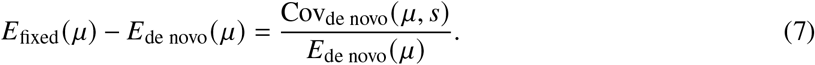

Equations 6 and 7 are again immediate consequences of Equations A2 and A5 in the mathematical Appendix.

### Correlation between mutation rates and selection coefficients

So far we have discussed the fact that the mean mutation rate and selection coefficient of mutations may differ depending on whether we consider random genetic perturbations (nominal distribution), random mutations introduced into a population (the *de novo* distribution) or random fixed mutations (the fixed distribution). However, our main interest here is in asking about whether there is a systematic relationship between mutation and selection, and how this relationship appears from each of these perspectives. We have already seen that such systematic relationships are important, as Equations 3 and 7 show how the covariance between mutation rates and selection coefficients in the nominal and *de novo* distributions determine how the means differ between these three joint distributions. Similarly, measures of association can sometimes have intuitive biological meanings in their own right. For example the sign of the correlation between mutation rates and selection coefficients in the nominal distribution determines whether the relationship between mutation rates and selection coefficients increases or decreases the expected rate of adaptive substitutions. This is because this correlation has the same sign as Cov_nom_(*μ, s*) = *E*_nom_(*μs*) − *E*_nom_(*μ*)*E*_nom_(*s*), which itself is simply the difference between a term proportional to the rate of adaptive substitutions, *E*_nom_(*μs*), and a second term proportional to what the rate of adaptive substitutions would be if mutation rates and selection coefficients were independently distributed, *E*_nom_(*μ*)*E*_nom_(*s*). Later we will see that the correlation between mutation rates and selection coefficients in the fixed distribution has a similar interpretation in terms of whether the relationship between mutation rates and selection coefficients increases or decreases the probability of parallel evolution.

Clearly the simplest possible case is when mutation and selection are completely independent. In particular, if *P*_nom_(*μ* = *μ*_*i*_ and *s* = *s*_*i*_) = *P*_nom_(*μ* = *μ*_*i*_)*P*_nom_(*s* = *s*_*i*_) for all *s*, then *P*_de novo_(*μ* = *μ*_*i*_ and *s* = *s*_*i*_) = (*μ*_*i*_ *P*_nom_(*μ* = *μ*_*i*_)/*E*_nom_(*μ*)) *P*_nom_(*s* = *s*_*i*_). Thus, if mutation and selection are independent with respect to the nominal distribution then they are also independent with respect to the *de novo* distribution. Moreover, the *de novo* distribution of selection coefficients remains unchanged relative to the nominal distribution and the *de novo* distribution of mutation rates is simply the size-biased version of the nominal distribution of mutation rates. Similarly, *P*_fixed_(*μ* = *μ*_*i*_ and *s* = *s*_*i*_) = (*μ*_*i*_ *P*_nom_(*μ* = *μ*_*i*_)/*E*_nom_(*μ*)) (*s*_*i*_ *P*_nom_(*s* = *s*_*i*_)/*E*_nom_(*s*)), so that the fixed distribution corresponds to independent draws from the sized-biased distribution of mutation rates and the size-biased distribution of selection coefficients. Thus, in the case of exact independence between mutation and selection, this independence is maintained in all three distributions. Importantly, as we shall see, exact independence is rarely realized when considering a specific finite set of mutations, which typically leads to substantial departures from these simple expectations.

As another simple scenario, consider only two possible mutation rates and two possible selection coefficients. For convenience, we will work with relative mutation rates and selection coefficients, so that we will write the lower mutation rate as 1 and the larger mutation rate as *B* > 1, and write the lower selection coefficient as 1 and the larger selection coefficient as *K* > 1. For this case it is also helpful to be able to work with the individual probabilities for the four possible combinations of mutation rate and selection coefficient. These probabilities are written as 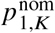, for example, referring to the probability of observing the lower mutation rate and the larger selection coefficient when drawing from the nominal distribution; likewise 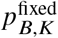 refers to the probability of observing the greater selection coefficient and greater mutation rate when drawing a random fixed mutation. For example, we will write 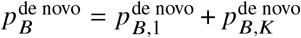 for the marginal frequency of the higher mutation rate in the *de novo* distribution and 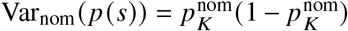 as the variance in the frequency of the high versus low selection coefficient under the nominal distribution. Finally,

Let 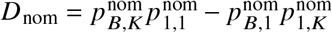, which is a measure of the association between the high and low levels for the mutation rate and selection coefficient.

With this basic setup in hand, we can now consider the Pearson’s correlation *r* between mutation rate and selection coefficient for our three distributions. In particular, we have

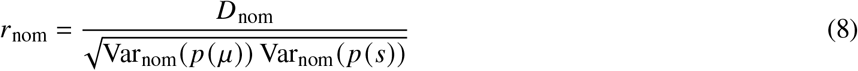

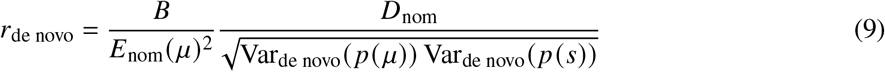

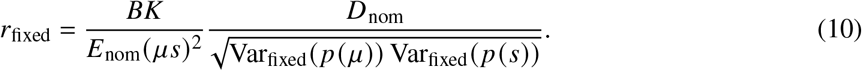

Note that *B, K, μ* and *s* are all positive, and variances are non-negative (and must be positive for the correlation to be defined), so that the sign of each expression depends only on *D*_nom_. Thus, we see that in the case where there are only two selective and two mutational classes, the sign of the correlation between mutation rates and selection coefficients is the same between all three distributions. This is illustrated in the first two rows of Fig. 2. In the first row, while the mean selection coefficient and mean mutation rate change across the three distributions, all three distributions remain uncorrelated. In the second row, the three distributions are all negatively correlated with the strength of the negative correlation increasing from nominal to *de novo* to fixed. Note that the negative correlation also results in a slightly non-monotonic pattern of change in the mean selection coefficient and mean mutation rate.

**Fig. 2.**
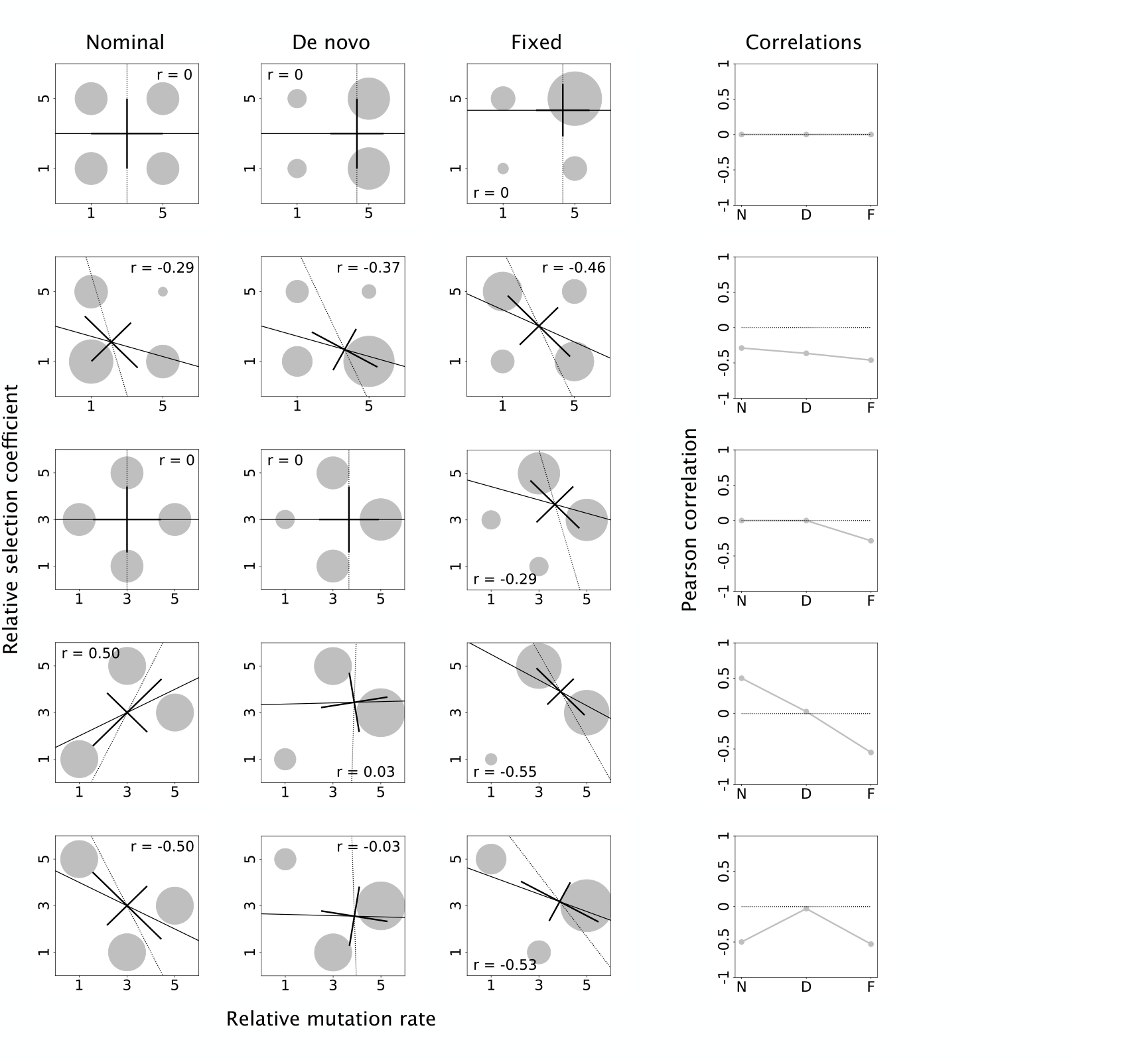
Correlations of mutation rates and selection coefficients under some simple models. Here mutation rates and selection coefficients each take on just 2 or 3 values. The first three columns show the nominal, *de novo*, and fixed distribution, with long lines giving the regressions of selection coefficient on mutation rate (solid) and mutation rate on selection coefficient (dotted). These lines intersect at the bivariate mean. The shorter lines are principal component axes with lengths corresponding to plus or minus one standard deviation. The fourth column shows how the Pearson’s correlation changes from nominal (N) to *de novo* (D) to fixed (F).

While the sign of the correlation of mutation rate and selection coefficient remains constant across all three distributions in the two-class case, if we extend the possibilities to 3 classes each of mutation rate and selection coefficient, the behavior of the correlation between mutation rates and selection coefficients becomes far less constrained. For example, the third row of Fig. 2 shows that mutation and selection can be uncorrelated in the nominal and *de novo* distributions but correlated among fixed mutations, unlike the case in row 1, which also has exactly 4 symmetric and equally sized *μ, s* categories.

More generally, we provide examples in Fig. S1 showing that, given just 3 distinct values for mutation rate and selection coefficient, any pattern of positive and negative signs is possible for the induced covariances. In addition, allowing mutation rates and selection coefficients to have different ranges of relative rates means that the signs of the correlations for the *de novo* and fixed distribution may depend on these specific ranges (Fig. S2).

What explains this counterintuitive behavior? The key observation here is that the lower moments of a size-biased distribution depend on higher moments of the original distribution. Thus, for example, the sign of the correlation between mutation rate and selection coefficient for the *de novo* distribution depends on one of the third mixed moments of *μ* and *s* in the nominal distribution, and specifically has the same sign as *E*_nom_(*μ* ^2^*s*)*E*_nom_(*μ*) − *E*_nom_(*μ* ^2^) *E*_nom_(*μs*) (this follows from Equation A7 in the Appendix, which gives the general formula for the variances and covariances of mutation rates and selection coefficients for all 3 joint distributions). Similarly, the fixed distribution depends on moments up to the fourth mixed moments of *μ* and *s*, and its correlation coefficient has the same sign as *E*_nom_(*μ* ^2^*s* ^2^)*E*_nom_(*μs*) − *E*_nom_(*μ* ^2^*s*)*E*_nom_(*μs* ^2^). Thus, many of the apparently paradoxical results above reflect this dependence on the higher mixed moments of the nominal distribution.

### Effects of a finite mutational target size

Our results above show that if the distributions of mutation rates and selection coefficients are independent for the nominal distribution then they are independent for all three distributions. However, in reality, the distributions are unlikely to be exactly independent, if only because the number of possible beneficial mutations in a given case is finite and may be quite small, e.g., exactly 9 beneficial options reported by Rokyta et al. (2005), or 11 beneficial variants that occurred at sufficient rates to measure both *s* and *μ* in Maclean et al. (2010). To illustrate the effects of having only a finite number of beneficial mutations, we consider the case where the underyling distributions of mutation rates and selection coefficients (that the realized possible beneficial mutations are sampled from) are independent and exponentially distributed, and we contrast our expectations for an infinite-sites model to the case where there are only a finite number of possible beneficial mutations.

The infinite sites case can be treated completely analytically. In particular, note that the size-biased version of an exponential distribution is a gamma distribution with the same rate and with a shape parameter of 2. Since a gamma distribution with an integer shape parameter *K* is the sum of *K* independent exponential distributions, we see that size-biasing the exponential is equivalent to taking the sum of 2 independent exponential random variables. Thus for an infinite-sites model with independent exponentially distributed mutation rates and selection coefficients, the mean mutation rate in the *de novo* distribution is twice that of the mean mutation rate in the nominal distribution, and the mean mutation rate and mean selection coefficient are both doubled in the fixed distribution relative to the nominal. More broadly, mutation rates and selection coefficient are independent random variables for all three of the nominal, *de novo*, and fixed distributions, where the *de novo* distribution has gamma-distributed mutation rates with a shape parameter of 2 and exponential selection coefficients, and the fixed distribution is gamma-distributed for both mutation rates and selection coefficients.

However, these simple results no longer hold if we relax the infinite sites assumption and consider a nominal distribution based on a finite sample from independent exponential distributions, as shown in Fig. 3. For instance, Fig. 3A shows a case in which the nominal distribution consists of 20 beneficial mutations, which are simulated as 20 pairs of *μ* and *s* values drawn independently from identical exponential distributions. From the nominal to the *de novo* to the fixed, the correlation shifts from mildly positive, to somewhat more positive, to substantially negative. In order to develop better intuitions for the typical size of the correlations that arise due to size-biasing a random sample and the variability in the signs of these correlations, in Fig. 3B, we have repeated this process 1000 times, for finite nominal distributions with 10, 100 or 1000 possible beneficial mutations.

**Fig. 3.**
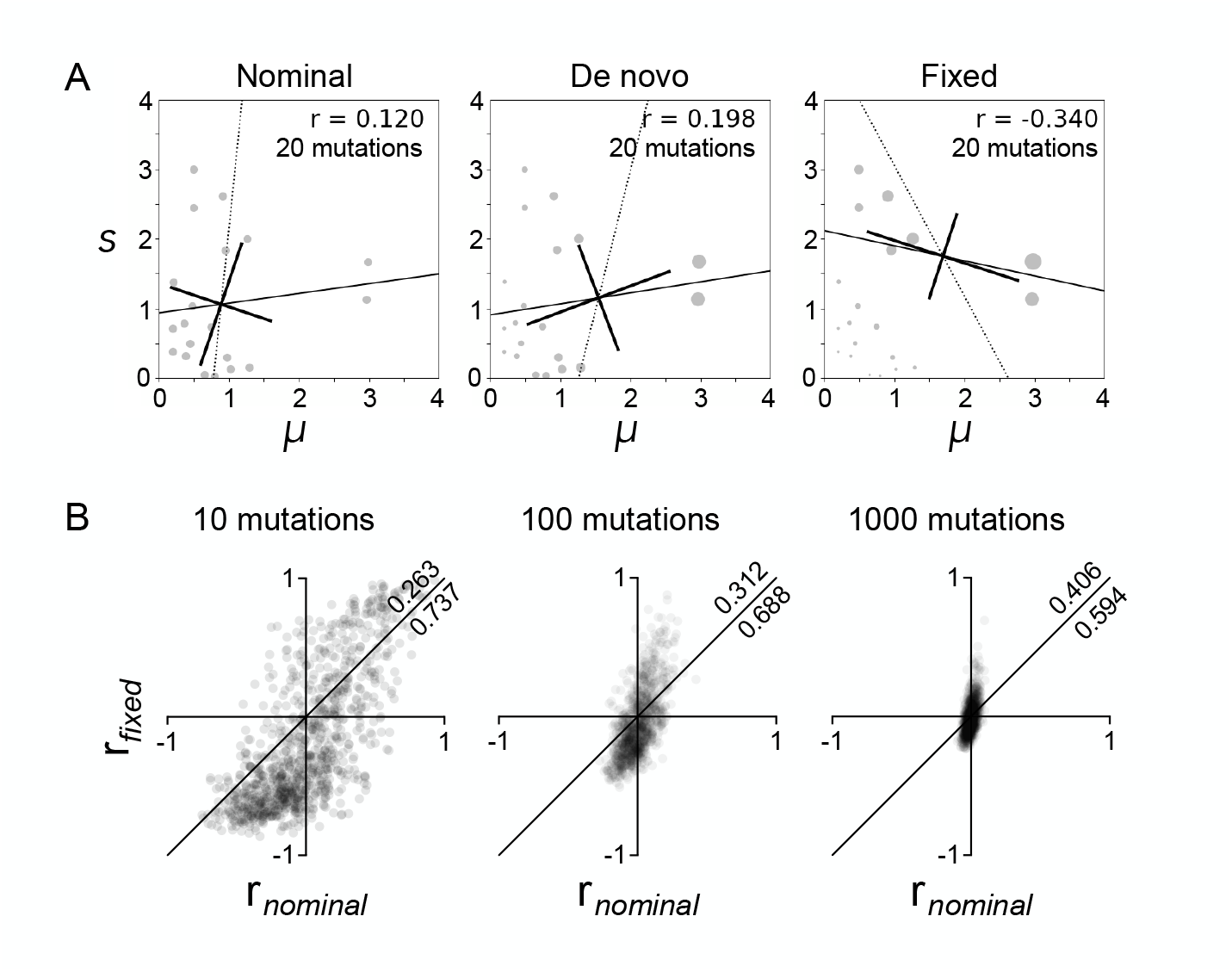
Effects of a finite mutational target size. **A** Here the fixed and *de novo* distributions are computed for a single example in which the nominal is a sample of 20 beneficial mutations with *μ*and *s* drawn independently from identical exponential distributions (each with a mean of 1). Regression lines and principal component axes are as in Fig. 2. An alternative example is given in Fig. S3A. **B** This same procedure is repeated 1000 times for samples of 10, 100, and 1000 beneficial mutations, revealing a tendency for the association to become more negative in the fixed distribution relative to the nominal distribution. Specifically, in bivariate scatterplots of *r* (Pearson’s correlation) for fixed vs. nominal, the fraction of points below the diagonal line of equality tends to be larger—sometimes by more than 2-fold—than the fraction above it. Comparable plots of *r* for fixed vs. *de novo* and *de novo* vs. nominal are shown in Fig. S3B.

While for increasingly large mutational targets the magnitude of the correlation between selection coefficients and mutation rates decreases (as we would expect, since the sample correlation coefficient is a consistent estimator of the underlying correlation), we see that the Pearson’s correlations often differ substantially from zero with 1000 beneficial mutations, and may become large with 10 or 100 beneficial mutations. Moreover, we observe a tendency for the fixed distribution to have a more negative correlation coefficient than the nominal distribution, i.e., the point (*r*_nom_, *r*_fixed_) tends to fall below the *y* = *x* diagonal shown in Fig. 3B (the fractions above and below this diagonal in each plot are given in the upper right corners, above and below the *y* = *x* diagonal respectively). Thus, for example, for a small mutational target size of 10 beneficial mutations, we observe a more negative correlation for the fixed distribution relative to the nominal distribution 74% of the time, and this value falls slightly to 69% for a moderately sized mutational target of 100 beneficial mutations.

While the induced correlations shown in Fig. 3 most frequently result in a more negative correlation in the fixed than the nominal, we also see a minority of samples showing strong positive correlations, where the magnitude of these positive correlations is larger than the typical magnitude of a negative correlation (Fig. 3B). Intuitively, both patterns arise because the correlation coefficient is strongly influenced by whether or not the set of beneficial mutations includes an outlier with both a large mutation rate and large selection coefficient. In most samples there is no such mutation, i.e., no density in the upper right of Fig. 3A, and this tends to produce a negative correlation in the fixed distribution. However, in the minority of samples when such a mutation is included by chance, this outlier mutation is strongly up-weighted in the fixed distribution, so that the correlation is dominated by the comparison between this outlier and the bulk of the other beneficial mutations, resulting in a positive correlation (for an example, see Fig. S3A).

More generally, this relationship between the shape of the nominal and the sign of an induced correlation in the fixed distribution provides intuitions for what to expect in natural systems, at least when there is no prior biological reason to suspect the nominal to show a systematic association of *s* and *μ* for beneficial mutations. Because natural distributions for both mutation rates and beneficial selection coefficients are typically L-shaped or positively skewed (i.e., few mutational hotspots, and few strongly beneficial mutations separate from the bulk of beneficial mutations near *s* = 0), the nominal distribution is unlikely to include a mutation with both an unusually large selection coefficient and an unusually high mutation rate. For such reasons, one may expect negative correlations in the fixed distribution to be more common, with occasional exceptions when the nominal happens to include a rare beneficial mutation favored both by a high mutation rate and a large fitness benefit.

### Non-parametric measures of association

The above results express the relationship between mutation rates and selection coefficients based on the Pearson’s correlation, which is a summary of the strength of the linear relationship between mutation rates and selection coefficients. While the Pearson’s correlation is a natural measure of association in this context due to the importance of the covariance in determining (for instance) the change in the mean selection coefficient between the nominal and *de novo* distributions, it is also reasonable to consider other forms of association between mutation rate and selection coefficient.

One such alternative approach is to ask about whether there is a systematic relationship between the rank ordering of mutation rates and selection coefficients. That is, if one draws two mutations with different mutation rates and selection coefficients, what is the probability that the mutation with the lower mutation rate also has the lower selection coefficient? Taking this probability of concordance between mutation rates and selection coefficients and subtracting the corresponding probability of discordance results in a non-parametric measure of correlation known as Goodman-Kruskal’s *γ* (Goodman and Kruskal, 1954), which takes values between +1 and -1 and can thus be directly compared to our results for the Pearson’s correlation. Specifically, if we take two random draws from the nominal distribution with selection coefficients *s*^(1)^ and *s*^(2)^, and mutation rates *μ*^(1)^ and *μ*^(2)^, and define *P*_nom_(concordance) = *P*_nom_(*s*^(1)^ < *s*^(2)^ and *μ*^(1)^ < *μ*^(2)^)+ *P*_nom_(*s*^(2)^ < *s*^(1)^ and *μ*^(2)^ < *μ*^(1)^) and *P*_nom_(discordance) = *P*_nom_(*s*^(1)^ < *s*^(2)^ and *μ*^1)^ > *μ*^(2)^)+ *P*_nom_(*s*^(2)^ < *s*^(1)^ and *μ*^(2)^ > *μ*^(1)^), then *γ* can be calculated as

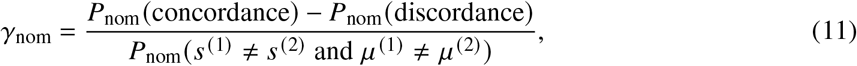

where we define *γ*_de novo_ and *γ*_fixed_ similarly and the denominator arises because we are conditioning on the two mutation rates and selection coefficients being distinct.

It is useful to compare the results for Goodman-Kruskal’s *γ* to the results we developed above for the Pearson’s correlation. In terms of results for simple theoretical distributions, it is easy to show that *γ*_nom_ = *γ*_de novo_ = *γ*_fixed_ = 0 if mutation rates and selection coefficients are independently distributed for any of the three distributions. Moreover, for the simple case with only 2 distinct mutation rates and selection coefficients it is also easy to show that *γ*_nom_ = *γ*_de novo_ = *γ*_fixed_, so that not only are the signs of the three correlations the same (as was the case for Pearson’s *r*), but the three coefficients are in fact identical (see Equations A20 and A22 in the Appendix). While these simple results suggest the intuition that the association between mutation rates and selection coefficients should be similar across all three distributions, this intuition is again strongly misleading. In more complex examples, Goodman-Kruskal’s *γ* (like the Pearson’s correlation) can take any sign pattern across the three distributions, as shown in Supplemental Fig. S4. Similarly, in Supplemental Fig. S5 we show that (1) finite mutational target size can also induce substantial changes in the magnitude and sign of *γ* across the three distributions, even when the mutation rates and selection coefficients are independent in the underlying distribution that this finite mutational target is drawn from, and (2) in the case where the underlying marginal distributions for mutation rates and selection coefficients are exponential, this finite sample effect tends to result in *γ*_fixed_ < *γ*_nom_, similar to what we observed for Pearson’s *r*.

## Empirical examples

As noted above, in principle mutation rates and selection coefficients may show a wide variety of strengths and patterns of association across the nominal, *de novo*, and fixed distributions. Therefore, it is of interest to consider the patterns of association observed in natural joint distributions. Here we consider a joint nominal distribution of selection coefficients and mutation rates reported for Dengue virus (Dolan et al., 2021), as well as data pertaining to the nominal and fixed distributions for mutations to TP53 in human cancers (Giacomelli et al., 2018).

### A nominal distribution for the Dengue virus genome

Dolan et al. (2021) carried out serial passages of Dengue virus on human host cells, and then used a high-accuracy deep-sequencing method to estimate the fitnesses of all possible 32,166 single-nucleotide variants across the entire Dengue genome. The same study provides estimates for the mutation rates of 12 types of nucleotide substitutions, uncovering the presence of a strong mutation bias towards *C* →*T*) transitions (note that, because the Dengue virus genome is single-stranded, a general nucleotide substitution model has 12 rates instead of the usual 6). We identified 237 single point mutations that had 95% confidence intervals for relative fitness that exceed 1 (using the high-confidence dataset of Dolan et al. 2021).

A substantial fraction of these mutations were strongly beneficial, with the largest estimated selection coefficient being 24. While such large *s* values are unusual in studies of population genetics, and they carry considerable uncertainty, very large selection coefficients are expected in this context given that (1) direct measurement of total population fitness showed an increase of 1 to 2 orders of magnitude (see Fig. 1 of Dolan et al. 2021), and (2) these dramatic increases were not due to a succession of fixed mutations with incremental effects, but to partial increases in frequency of multiple large-effect mutations. Many-fold increases in fitness are biologically plausible when infectivity on host cells in culture is low and can increase greatly, as is the case for some of these mutations (Dolan et al., 2021). However, because these large selection coefficients violate our assumption that the probability of fixation is proportional to *s*, in what follows we will consider both a direct application of our previous theory—which is like assuming that the reported *s* values can be rescaled to smaller absolute values—and simulations that capture the saturation of the probability of fixation (for large *s*) as well as clonal interference between multiple adaptive lineages. More generally, our goal here is not to attempt to replicate the actual population biology of Dengue virus, which is complex: we are merely taking advantage of an empirical, genome-wide nominal distribution that has been measured in Dengue virus for a single environment, and considering the consequences of this distribution in light of our analytical theory and in the context of relatively simple population-genetic simulations.

Importantly, while the distribution of beneficial selection coefficients ranges over several orders of magnitude, so do the observed mutation rates (Fig. 4A), and there are strong outliers in terms of both selective and mutational advantage. In particular, the estimated rates of *C* →*T*) transitions is 22 times larger than the next highest mutation rate. Overall, there appears to be a weak negative association between mutation rates and selection coefficients in the nominal distribution, with a Goodman-Kruskal’s *γ* of −0.16 (*P* < 10^−6^) and a statistically non-significant Pearson’s *r* of −0.06 (indicating that we cannot reject the hypothesis that the underlying distribution of mutation rates and selection coefficients is uncorrelated).

**Fig. 4.**
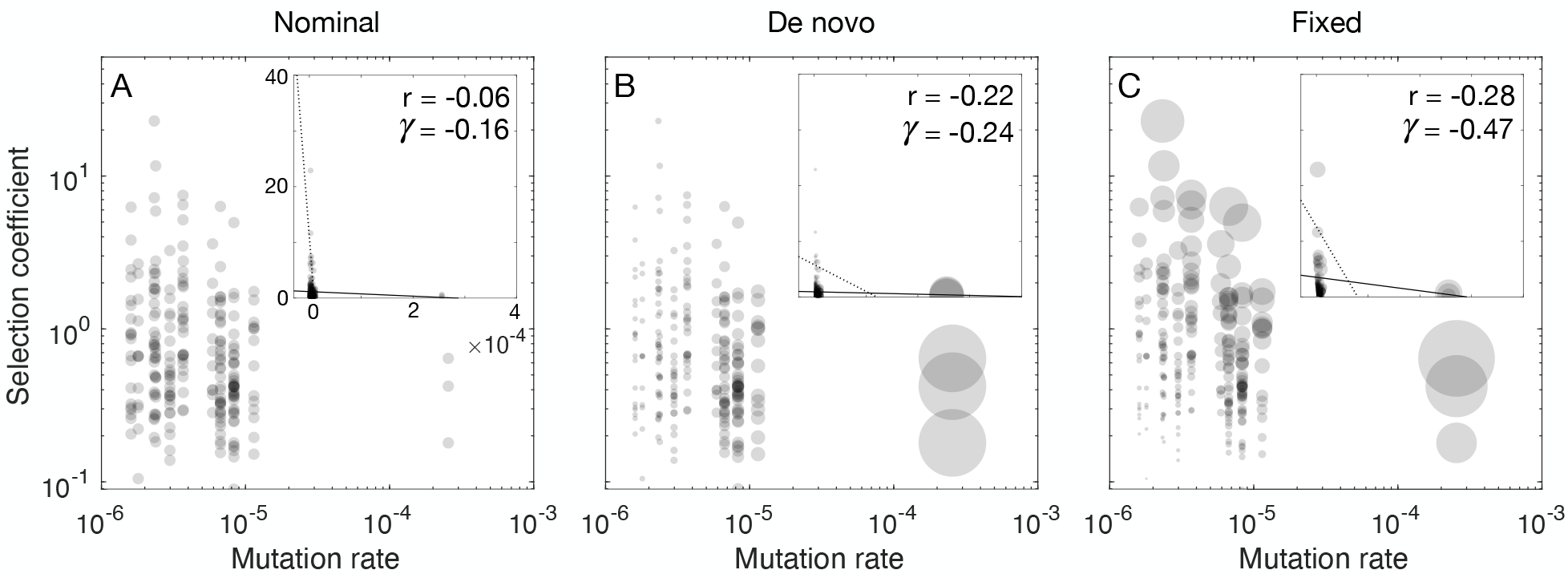
Calculated effects on an empirical nominal distribution. **A**. The nominal distribution consists of 237 beneficial point mutations reported for Dengue virus by Dolan et al. (2021). The inset shows the distribution on a linear scale, including the regression of *s* on *μ* (solid line) and *μ*on *s* (dotted line). Pearson’s correlation coefficient *r* = −0.06 (*P* = 0.28), Goodman-Kruskal’s *γ* = −0.16 (*P* < 10^−6^). **B**. The *de novo* distribution shows the nominal distribution weighted by the mutation rates; A = −0.22 (*P* < 10^−6^) and *γ* = −0.24 (*P* < 10^−6^). **C**. The fixed distribution shows the *de novo* distribution weighted by the selection coefficients; *r* = −0.28 (*P* < 10^−6^) and *γ* = −0.47 (*P* < 10^−6^). The mean values are: 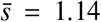 and 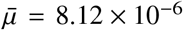 (nominal), 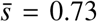 and 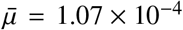 (*de novo*), and 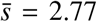 and 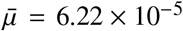 (fixed).

However, this negative association is amplified in the *de novo* distribution, largely due to the very high rate of occurrence of the *C* →*T*) transitions (the 3 beneficial *C* →*T*) transitions make up ∼ 40% of all *de novo* mutations), and this results in a Pearson’s correlation coefficient of *r* = −0.22 (*P* < 10^−6^), and a Goodman-Kruskal’s *γ* of -0.24 (*P* < 10^−6^). Thus, a slight negative association of mutation rates and selection coefficients among distinct advantageous mutations is transformed into a substantial negative association among *de novo* mutations. The negative association also results in a substantial decrease in the mean selection coefficient from 1.14 in the nominal distribution to 0.73 in the *de novo* distribution.

Finally, we can consider the predicted joint distribution of mutation rates and selection coefficients for fixed mutations. Applying our analytical theory to the *de novo* distribution results in *r* = −0.28 (*P* < 10^−6^) and Goodman-Kruskal’s *γ* = −0.47 (*P* < 10^−6^), indicating a moderate negative association that reflects both the fixation of mildly beneficial changes favored by mutation (e.g. ∼ 23% of the total fixed mutations are the three beneficial *C* to *T* transitions) and an enrichment for low-rate mutations with large selection coefficients (e.g. the average mutation rate is 72% larger among *de novo* mutations than it is among fixed mutations).

Whereas Fig. 4C shows a fixed distribution based on linear size-biasing (i.e., multiplying by *s*), in reality we might expect departures from this simple theory due to the fact that the survival probability for a new mutation saturates near its maximum of 1 for large *s*, and due to competition between multiple beneficial mutations, i.e., clonal interference (Gerrish and Lenski, 1998). To address these additional factors we also conducted population simulations using SLiM v3.4 (Messer, 2013) to estimate the fixed distribution. For each run of the simulation, we recorded the identity of all adaptive mutations on the first sequence to reach fixation, repeating this process until we obtain 1500 beneficial substitutions. In order to consider a wide range of population-genetic conditions, we let the population size vary over 4 orders of magnitude. This produced a wide degree of variation in the expected number of beneficial mutations per generation, which we write as *Nμ*_tot_ (population size times the total beneficial mutation rate).

The results of these simulations are shown in Fig. 5. The resulting joint distribution for the mutation rates and selection coefficients of fixed mutations depends on mutation supply (*Nμ*_tot_), as shown by the examples in Fig. 5A and 5B, which represent the lowest and highest values for mutation supply, respectively. For low values of mutation supply, the fixed distribution (Fig. 5A) is similar to the distribution based on linear enrichment by *s* (Fig. 4C) but with less of an enrichment for large selection coefficients. However, for large values of mutation supply (Fig. 5B) we see a considerable decrease in the number of fixations of high-rate *C*→*T* transitions and instead see an enrichment for large selection coefficients. This results in an overall decrease in the magnitude of the negative correlation between mutation rates and selection coefficients among fixed mutations (Fig. 5C), but a relatively constant value for Goodman-Kruskal’s *γ* (Fig. S6). Intuitively, this difference between *γ* and *r* arises because the mutations whose contributions to the fixed distribution change the most as a function of mutation supply are the high-*μ*, low-*s* outliers and the low-*μ*, high-*s* outliers, and these outliers have much more leverage on *r* than *γ* because the latter is non-parametric, and thus reflects only the ranks and not the distance of these outliers to the bulk of the other mutations.

**Fig. 5.**
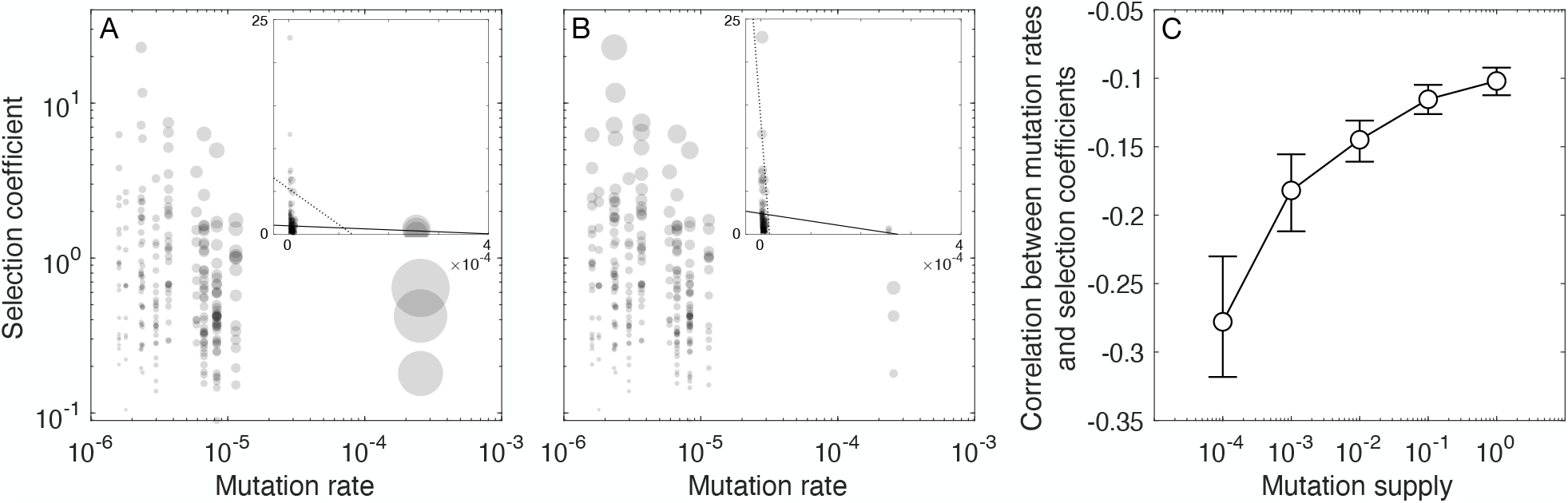
Mutational supply shapes induced associations. Panel **A** shows simulations with a mutation supply of *Nμ*_tot_ = 10^−4^, giving *r* = −0.28 (*P* < 10^−6^). Mean values are 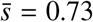 and 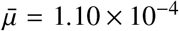. For panel **B**, *Nμ*_tot_ = 10^0^ and *r* = −0.10 (*P* = 0.005), and mean values are 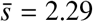 and 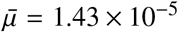. The insets show a linear scale, while the log-scale clarifies that the bulk of mutations have both low *μ* and small *s* (*r* is for unlogged values). Panel **C** shows the correlation for the fixed distribution as a function of beneficial mutation supply (*Nμ*_tot_). Here # ranges from 12 to 1.2 × 10^5^. Error bars depict 95% bootstrap confidence intervals based on 10^3^ bootstrap samples.

### Nominal and fixed distributions for TP53 mutations in human cancer

In the previous section we used empirical data to infer the nominal distribution of possible mutations, which we used to derive predictions for the fixed distribution using simple population-genetic models and computer simulations. However, one important limitation of that dataset is the lack of information about the actual set of mutations that reached fixation in nature, which is why we turned to evolutionary simulations.

In the case of somatic mutations in cancer, useful data are available relating both to the nominal distribution, and to the distribution of variants after mutation and somatic expansion, i.e., cancerous growth. One example of a protein for which such data are available is p53, encoded by the TP53 gene—the most frequently mutated gene in many forms of human cancer, sometimes called “the guardian of the genome” for its role in conferring genetic stability and preventing both genome mutation and cancer formation (Hainaut and Pfeifer, 2016). Using deep mutational scanning, Giacomelli et al. (2018) generated all 7880 possible non-synonymous amino acid changes to the p53 protein and measured the growth rate that each variant conferred on human lung carcinoma cells in the presence and absence of endogenous p53. As with the previous dataset, we focus only on those amino acid changes that confer a selective advantage (combined enrichment score *R*_all_ > 0, see Methods). In the context of studying cancer, a selective advantage refers to somatic growth rate greater than that conferred by the wild-type sequence, and thus a large fraction (31%) of changes are advantageous (i.e., beneficial for tumor growth) due to loss of the protective function of p53. As explained in Methods, selection coefficients from different p53 deep mutational scanning studies (Giacomelli et al., 2018; Kotler et al., 2018; Staller et al., 2022) correlate well (Carbonnier et al., 2020), but the absolute scale of the selective advantage is uncertain. We therefore consider three different scales: the unusually high *s* values based on Giacomelli et al. (2018), the same values divided by 10, and the same values divided by 100. Note that changing the scale of *s* does not affect calculated correlations based on size-biasing, but may strongly influence the relative chances of fixation in population-genetic simulations.

To empirically characterize the relative mutation rates of each possible amino acid change, we used data from the Pancancer Analysis of Whole Genomes database (Weinstein et al., 2013). We estimated the relative mutation rates using a trinucleotide context model, so that the rates of each nucleotide change are specified given the identity of the bases that immediately flank the mutated base (Methods, Fig. S7). We then calculated the estimated relative mutation rate for each amino acid change as the sum of the rates for each of its associated single nucleotide changes. Note that these are relative rates: the absolute scale is not known (see Methods).

Here, we combine these empirically estimated mutation rates and selection coefficients to construct a nominal distribution that contains mutation rates and selection coefficients for all 1489 possible beneficial amino acid changes (Fig. 6A). The shape of the nominal distribution for TP53 is different than the one for Dengue virus (Fig. 5A), particularly with regard to the presence of beneficial mutations that have both high selection coefficients and high mutation rates, which were absent in the Dengue virus dataset. While the correlation between mutation rates and selection coefficients in the nominal distribution is similar to that for Dengue virus, being close to zero and slightly negative (Pearson’s correlation coefficient *r* = −0.04 with *P* = 0.03, and *γ* = −0.09 with *P* < 10^−6^), the transformations to the *de novo* distribution (Fig. 6B) and the fixed distribution based on linear size-biasing (Fig. 6C) are quite different. In particular the *de novo* distribution is almost uncorrelated, while the fixed distribution becomes slightly positively correlated (*r* = 0.10 with *P* < 10^−^6, Goodman-Kruskal’s *γ* = 0.04, *P* < 10^−6^). These results are consistent with the intuition that the presence of mutations with large mutation rates and also large selection coefficients will tend to promote a positive correlation between *μ* and *s* among fixed mutations.

**Fig. 6.**
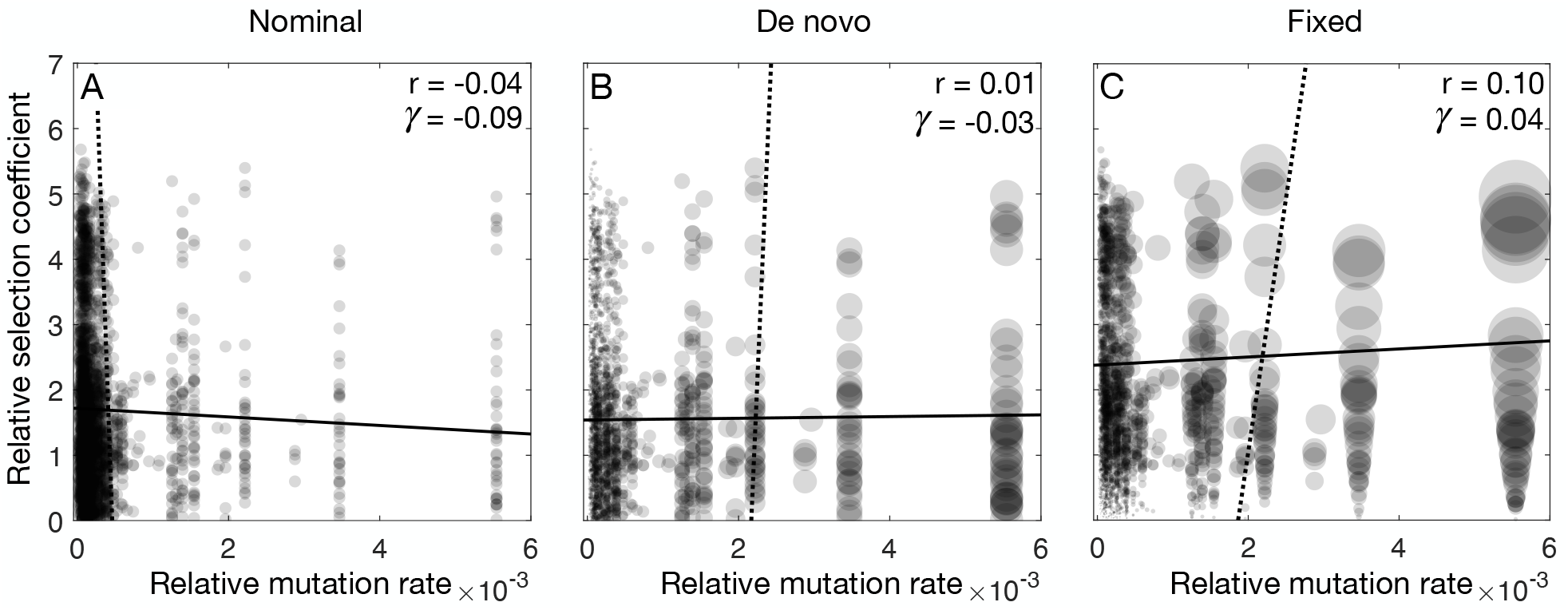
Calculated distributions from an empirical nominal distribution for TP53 mutations. **A**. The nominal distribution shows 1489 single-nucleotide beneficial amino acid changes with selection coefficients estimated from deep mutational scanning and relative mutation rates from a trinucleotide context model. Pearson’s correlation coefficient *r* = −0.04 (*P* = 0.03), Goodman-Kruskal’s *γ* = −0.09 (*P* < 10^−6^). Mutation rates are expressed as the estimated fraction of mutations in TP53 of each type. **B**. The *de novo* distribution, reflecting size-biasing by mutation rates. Pearson’s correlation coefficient *r* = 0.01 (*P* = 0.8), Goodman-Kruskal’s *γ* = −0.03 (*P* < 10^−6^). **C**. The fixed distribution shows the *de novo* distribution weighted by the selection coefficients. Pearson’s correlation coefficient *r* = 0.10 (*P* < 10^−6^), Goodman-Kruskal’s *γ* = 0.04 (*P* < 10^−6^). The mean selection coefficient 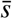 and mean mutation rate 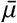 for each distribution are: 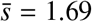 and 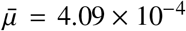 for the nominal, 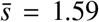 and 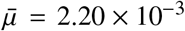 for the *de novo*, and 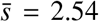 and 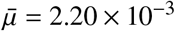 for the fixed distribution.

The analog of the fixed distribution for TP53 variants after mutation and selection is the distribution of mutations in clinical samples from cancer patients. Here we use frequency data from the GENIE database of the American Association for Cancer Research (AACR Project Genie Consortium et al., 2017), allowing for the assignment of observed frequencies to each possible beneficial amino acid change identified in the nominal distribution (Methods). The distribution of fixed mutations shown in Fig. 7A reveals an even stronger induced positive correlation than expected from the linear enrichment used above (*r* = 0.22 with *P* < 10^−6^; *γ* = 0.08 with *P* < 10^−6^), as well as an increased mean selection coefficient among fixed mutations of 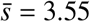. Because one explanation for this increased enrichment for large selection coefficients is competition between beneficial mutations due to clonal interference (Gerrish and Lenski, 1998), we again conducted population simulations with a variable mutation supply *Nμ*_tot_ (see Methods). We find a much closer fit with the observed data at an intermediate mutation supply (Fig. 7B). However, the correlation between mutation rates and selection coefficients is still only approximately 0.1 for both *r* and *γ*, considerably lower than the value of 0.22 seen in the observed TP53 data. In general, if the estimated mutation rates and selection coefficients were fully accurate, the observed frequencies in the fixed distribution should be continuous functions of the mutation rate and selection coefficient (plus Poisson-distributed noise in the numbers of clinical observations), which is in contrast to the heterogeneity we observe in Fig. 7A. Thus, while the pattern of observed mutations supports the relevance of our mutation rate and selection coefficient estimates to clinical data, it also suggests the presence of additional unaccounted factors such as mutational heterogeneity not captured in the trinucleotide model, or discrepancies between the laboratory estimates of selection coefficients and the selection coefficients that are relevant during somatic evolution.

**Fig. 7.**
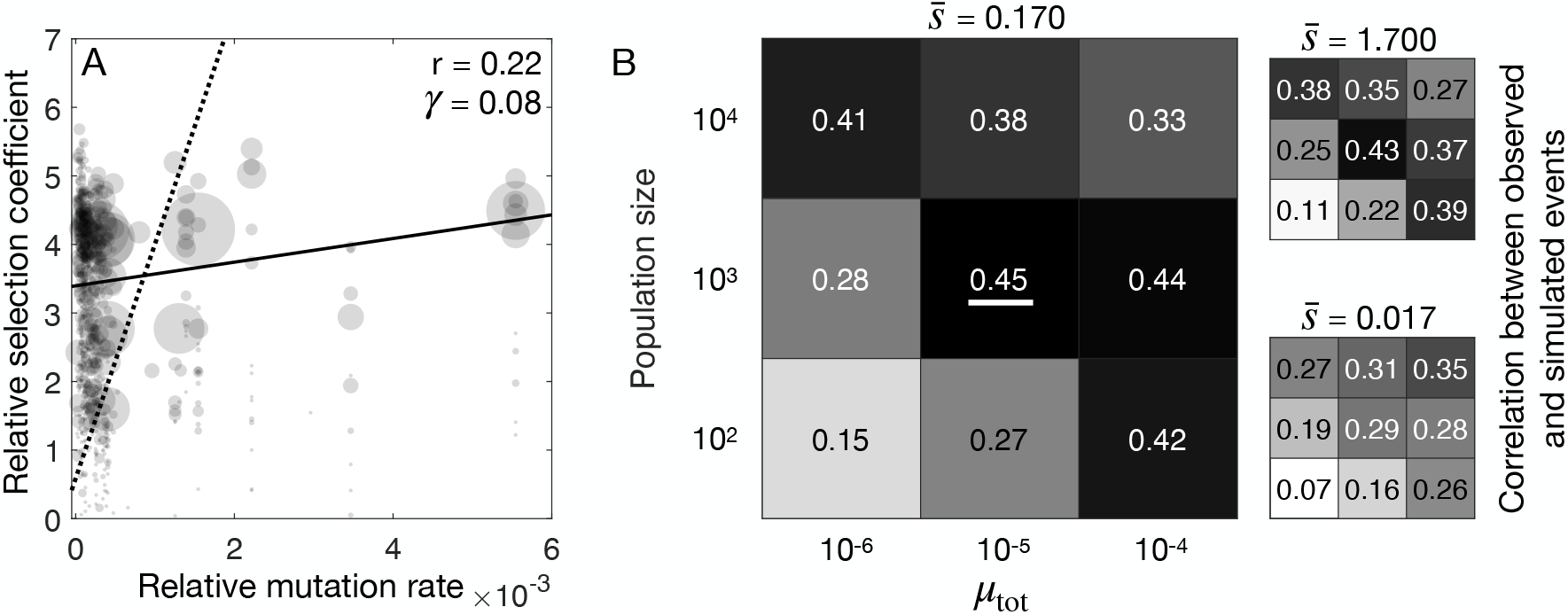
Population genetic conditions shape the distribution of fixed adaptive mutations. **A**. To represent the observed fixed distribution for the TP53 case, relative mutation rates and selection coefficients are weighted by the observed frequency of mutations in the GENIE database. The mean relative selection coefficient is 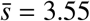 and the mean relative mutation rate is 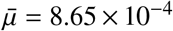. Pearson’s correlation coefficient *r* = 0.22 (*P* < 10^−6^), Goodman-Kruskal’s *γ* = 0.08 (*P* < 10^−6^). **B**. Results of population simulations of the fixed distribution, showing how the degree of correlation between the observed and simulated versions of the fixed distribution depends on *N, μ*_tot_ and the scaling of relative selection coefficients. Each 3 × 3 grid shows the correlation coefficients *r* for 9 combinations of *N* and *μ*_tot_, for relative selection coefficients reduced 10-fold (left, 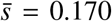), unscaled (right, upper), and reduced 100-fold (right, lower). The 95% bootstrap confidence intervals are given in Table S1.

Finally, we wished to probe the effect of mutation rate further by considering multi-nucleotide mutations as an additional class of mutations with rates much lower than the single-nucleotide mutations treated above. While multi-nucleotide mutations are often ignored in models, they are known to occur widely in nature, at a combined rate roughly 2 or 3 orders of magnitude lower than the combined rate of single-nucleotide mutations (Schrider et al., 2011; Smith et al., 2003). Thus one may imagine that multi-nucleotide variants frequent in clinical studies must have had strong selective effects to compensate for their low mutation rates, particularly in a clonal interference regime with more frequently arising single-nucleotide mutations.

The study of TP53 by Giacomelli et al. (2018) mentioned above, like most deep mutational scanning studies, covers all amino acid changes, not just the 150 (out of 380) types of replacements possible via single-nucleotide mutations, thus it provides a nominal distribution of selection coefficients for both the single- and multi-nucleotide variants. In addition, the data on cancer prevalence provides the fixed distribution for multi-nucleotide variants. We do not have a detailed empirical distribution of mutation rates for multi-nucleotide mutations. However, in the absence of a detailed rate model, we can simply compare two mutational categories and apply rank-order statistics to the fitness distributions. If two values are drawn from the same distribution, the chance that the first one is greater is 50%; likewise, the null expectation that a randomly chosen multi-nucleotide variant is fitter than a random single-nucleotide variant is 50%, and the Mann-Whitney U test tells us how significant is any deviation from this value.

The results of this comparison are shown in Fig. 8. The nominal distribution for beneficial changes consists of 3204 multi- and 1489 single-nucleotide variants (a ratio of 2.15, slightly lower than the ratio of 2.42 in the entire DFE). The multi-nucleotide variants have a slight advantage, namely a multi-nucleotide variant has a ∼ 61% chance of being fitter than a single-nucleotide variant (Mann-Whitney U test, *U*/(*n*_1_ * *n*_2_) = 0.61, *P* < 10^−6^, 95% CI, 0.59 to 0.62). This difference is magnified considerably in the fixed distribution (i.e., clinical prevalence). In the fixed distribution, multi-nucleotide variants are quite rare, reflecting their low mutation rates, but a random multi-nucleotide variant is fitter than a random single-nucleotide variant ∼ 88% of the time (Mann-Whitney U test, *U*/(*n*_1_ * *n*_2_) = 0.88, *P* < 10^−6^, 95 *P* CI, 0.82 to 0.94), significantly greater than in the nominal (95% bootstrap confidence intervals for the normalized Mann-Whitney test statistic *U*/(*n*_1_ * *n*_2_) based on 10^3^ bootstrap samples do not overlap). The mean selection coefficient 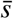 for multiples and singles changes from 2.14 (95% CI, 2.08 to 2.19) and 1.48 (95% CI, 1.42 to 1.54) in the nominal, to 4.15 (95% CI, 3.77 to 4.56) and 2.33 (95% CI, 2.24 to 2.43), respectively, in the fixed distribution. We additionally test the robustness of our results by once again employing Goodman-Kruskal’s *γ* to characterize the association between mutation rates and selection coefficients. This statistic shows a similar trend to the Mann-Whitney U test: for the nominal distribution of beneficial changes *γ* = −0.21 (*P* < 10^−6^), whereas for the fixed distribution of beneficial changes *γ* = −0.76 (*P* < 10^−6^). Thus, among TP53 mutations sampled from human tumors, there is a strong tendency for the multi-nucleotide mutations to have greater fitness as measured in the Giacomelli et al. (2018) laboratory experiments.

**Fig. 8.**
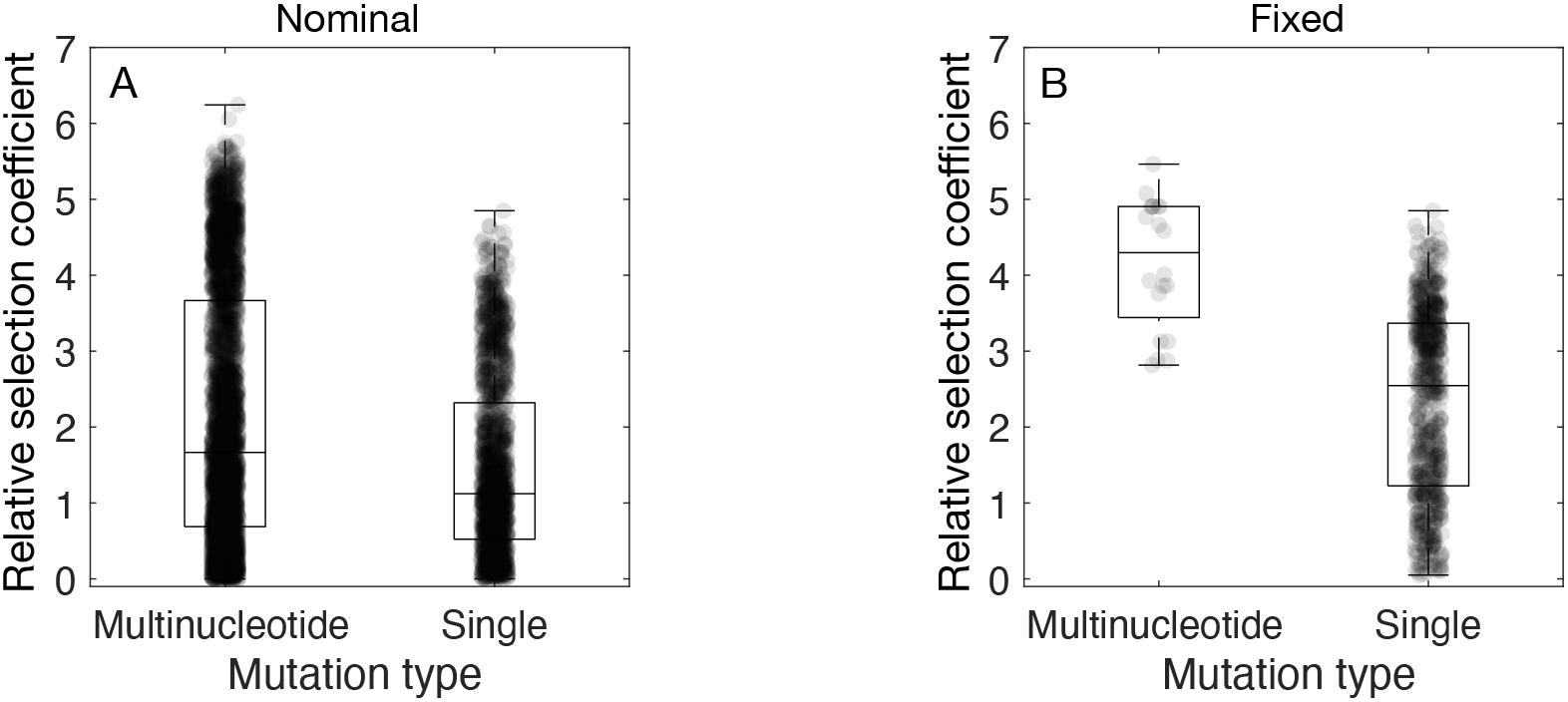
Fixed multinucleotide mutations have stronger fitness effects. Amino acid changes are aggregated into two categories, depending on whether they take place by a single-nucleotide or multi-nucleotide mutation. **A**. Nominal distribution of beneficial amino acid changes, showing a slight advantage of multi-nucleotide changes, with mean *s* of 2.1 vs. 1.5 for single-nucleotide changes. **B**. Observed distribution of fixed beneficial amino acid changes, showing a much stronger advantage of multi-nucleotide changes, with mean *s* of 4.2 vs. 2.3 for single-nucleotide changes.

## Discussion

The existing literature suggests a complex relationship between mutation and selection in determining the distribution of beneficial variants that contribute to adaptation (Cano et al., 2023). For instance, the most common outcomes of adaptation are often the ones that are highly mutationally likely, rather than the most beneficial, e.g., Giacomelli et al. (2018) report that 30% of TP53 tumor-associated mutations occur at 5 sites subject to heightened CpG mutations, and Cannataro et al. (2018) suggest that the most prevalent driver mutations are often not the ones with the highest growth rates, but high rates of mutation. Likewise Leighow et al. (2020) report that the most clinically common Imatinib-resistant variants among leukemia patients are not the ones that provide the greatest resistance, but the ones that arise at higher rates of mutation. Some results hint further that the less mutationally likely outcomes tend to have higher fitness benefits, and *vice versa*, e.g., Fig. 2 of Cannataro et al. (2018) appears to show a negative correlation between selection intensity and mutation rates (as does the comparable Fig. 3 of Cannataro et al. 2019). In Fig. 2A of Watson et al. (2020), the joint distribution of mutation rates and selection coefficients estimated for the 20 most common clonal haematopoesis mutations clearly shows a negative correlation. Stoltzfus and Norris (2016) report that, among studies that measure fitness of nucleotide changes in laboratory adaptation experiments, transversions (which happen at a lower rate) seem to have slightly larger fitness benefits than transitions.

Such results raise the question of what kinds of associations between mutation rates and selection coefficients are expected to emerge, whether they will be strong enough to observe, and whether they might tell us something important about the nature of the relevant adaptive process. Here we have developed an initial theory to address such questions, focusing on a regime of mutation-limited adaptation in which the distribution of fixed beneficial mutations is related to the distribution of possible beneficial mutations via a linear enrichment for mutations with high mutation rates and a linear enrichment for mutations with large selection coefficients.

Of interest is whether the process of adaptation induces associations between mutation rates and selection coefficients among fixed mutations, and whether these associations tend to be positive or negative. We find that, in theory, a variety of effects are possible. This makes the issue much more empirical, dependent on the shapes of joint distributions found in nature. Using available data from Dengue virus, we find that a simple population-genetic model of adaptation leads to a negative correlation between mutation rate and selection coefficient among fixed mutations, one that becomes stronger when mutation supply is low. The data available on TP53 mutations allows us to compare an observed fixed distribution (based on clinical prevalence) to a nominal distribution based on deep mutational scanning, and also to compare this observed fixed distribution to distributions obtained via population-genetic simulations. For single-nucleotide changes, we find a positive association of mutation rate and selection intensity in clinical prevalence data. Simulations from the nominal distribution also yield a mostly positive association. However, when we compare single-to multi-nucleotide variants, we observe a weak negative association in the nominal distribution and a much stronger negative association in the fixed distribution. That is, in clinical data based on the prevalence of mutations among patients, the multi-nucleotide changes, which emerge by mutation at a far lower rate, appear to be have considerably greater growth advantages than single-nucleotide changes.

These two empirical nominal distributions (for Dengue virus and TP53) exhibit qualitative and quantitative differences that explain the opposition in sign of the induced correlation between mutation rates and selection coefficients in the distribution of fixed mutations. In both distributions, *C* → *T*) transitions occur at the highest rate, but mutation bias in the Dengue dataset is much greater. In the nominal distribution of Dengue virus mutants, the mutationally favored *C* →*T*) transitions generally provide only modest selective advantages, generating an “L” shape reflecting the lack of options with both high mutation rate and high selection coefficient. Based on our analytical results, this shape of the nominal distribution will tend to induce a negative correlation in the fixed distribution. In contrast, in the case of TP53, due to both the lower mutation bias in the nominal distribution and to clonal interference, selection coefficients have a considerably stronger influence on the fixed distribution than mutation rates. Thus, the theoretical framework developed here, together with simulations, provides some useful guidance for understanding how associations between mutation rate and selection coefficient are influenced by the nominal distribution, its associated mutation biases, and population-genetic conditions.

These empirical arguments must be interpreted cautiously. Clearly, researchers conducting deep mutational scanning studies have struggled to improve measurements of fitness (and other functional effects) so that they are less noisy and less likely to be confounded with mutability (e.g., Acevedo et al. 2014). Likewise, mutation-accumulation studies are designed to remove effects of selection, so as to accurately measure mutational spectra (Katju and Bergthorsson, 2019). However, our analysis here, focusing specifically on correlations, subjects these data to a much higher level of scrutiny of the joint distribution than was imagined for the original uses of the data. With clinical data on cancer, for instance, there is no clear quantitative standard of ascertainment that specifies what qualifies as rapidly growing cancerous tissue. Ultimately, our understanding of what types of associations are induced in nature may require new methods designed specifically to characterize these joint distributions of mutation rates and selection coefficients without bias.

With these caveats in mind, the results reported here suggest the following: (1) the dual dependency of adaptation can induce strong associations between mutation rates and selection coefficients among fixed mutations; (2) the sign of the association between mutation rates and selection coefficients can change across the three joint distributions, based on higher moments of the nominal distribution; (3) weak higher-order associations (in the nominal) due to finite sampling may be amplified into substantial associations among fixed mutations; and (4) in practice, according to limited evidence currently available, all of these theoretical points may be highly relevant to interpreting natural cases.

This work also highlights the importance of distinguishing different distributions of mutational effects. Mutation accumulation studies typically sample from a *de novo* distribution in which mutations with higher rates are observed more frequently. By contrast, mutation-scanning studies typically are designed to draw from a nominal distribution without bias, using engineered variants. The term “DFE” is used in the empirical literature to refer to all three kinds of distributions of mutational effects, including nominal distributions from deep mutational scanning studies, *de novo* distributions from mutation accumulation, and occasionally for fixed distributions. Studies of theoretical population genetics for asexuals most often use the *de novo* distribution, e.g. in the notation of Good et al. (2012), *ρ*(*s*) is the *de novo* distribution of selection coefficients, and *ρ*_*f*_ (*s*) is the fixed.

A further source of complexity is that the nominal—which determines what is included in the other two distributions—does not have a single definition, as a matter of principle. Mutation samples from the possible, and selection samples from the actual. But there is no *a priori* definition of what is possible, therefore the nominal distribution has no *a priori* definition. For example, Sanjuan et al. (2004) measure beneficial variants among single-nucleotide mutations by engineering a specific list of changes; Wu et al. (2014) sample from this distribution in a somewhat biased way using error-prone PCR; other deep mutational scanning studies, e.g., Thyagarajan and Bloom (2014), randomize triplet codons in order to consider all single-amino-acid variants, rather than all single-nucleotide variants. Several studies have explored *combinations* of point mutations (e.g., Diss and Lehner 2018). Recently, Macdonald et al. (2022) reported a new deep mutational scanning method that covers small insertions and deletions, in addition to nucleotide substitutions. These more expansive ways of defining the nominal cover a greater range of potentially important types of mutations (i.e., beyond nucleotide substitutions), but they do not exhaust the universe of mutations, which is is astronomically large and dominated by complex events that occur at low rates (as explained in Appendix B of Stoltzfus 2021). In practice, the nominal distribution of beneficial mutations will often be defined by the observed *de novo* mutations that appear at a rate large enough and with a fitness effect large enough to be readily observable.

As noted above, the data used in this study were not designed to be used in this way and are not ideal. What kinds of systems and protocols would produce more appropriate data? Foremost, progress on this topic absolutely requires studies that measure *s, μ* and the frequency evolved *f*_*e*_ in a clearly defined system. Second, models of average rates for categories of mutations are a poor substitute for actual rates for each individual mutational change. For instance, Maclean et al. (2010) measured mutation rates for just 11 nucleotide substitution mutations in the same gene, finding a 30-fold range of rates. Hodgkinson and Eyre-Walker (2011) estimated that a triplet context model captures only 1/3 of the variance in individual rates of nucleotide substitution mutations. Third, the problem of estimation error requires explicit attention, particularly the possibility that, when *μ* and *s* are estimated together using a population-genetic approach (e.g., as in Watson et al. 2020), the errors may be correlated. An ideal system to collect such measurements would allow some flexibility to adjust various factors such as population size *N* (e.g., via droplet technology, as in Ruelens and de Visser 2021), the strength of selection (e.g., via concentration of antibiotic), and the size of the mutational target (e.g., aggregating results over a narrow or broad set of antibiotics).

Finally, let us consider three general directions for further theoretical work. First, while our focus here has been on the relationships of the nominal, *de novo* and fixed distributions, two other joint distributions, shown in Fig. 9, can also be derived by further size-biasing and warrant additional attention (see the mathematical Appendix for details). Specifically, weighting the fixed distribution again by selection coefficients gives a distribution that corresponds to the contribution of each possible beneficial mutation to the expected rate of adaptation (i.e., the expected rate of fitness increase). Size-biasing this distribution again with respect to mutation rates then produces a distribution that gives the contribution of each mutation to the overall probability of parallel adaptation, defined as the probability that the same mutation would be fixed twice in two independent bouts of adaptation (Chevin et al., 2010; Lenormand et al., 2016). Moreover, as shown in the Appendix, each of the normalizing constants for the 5 distributions also has an evolutionary interpretation.

**Fig. 9.**
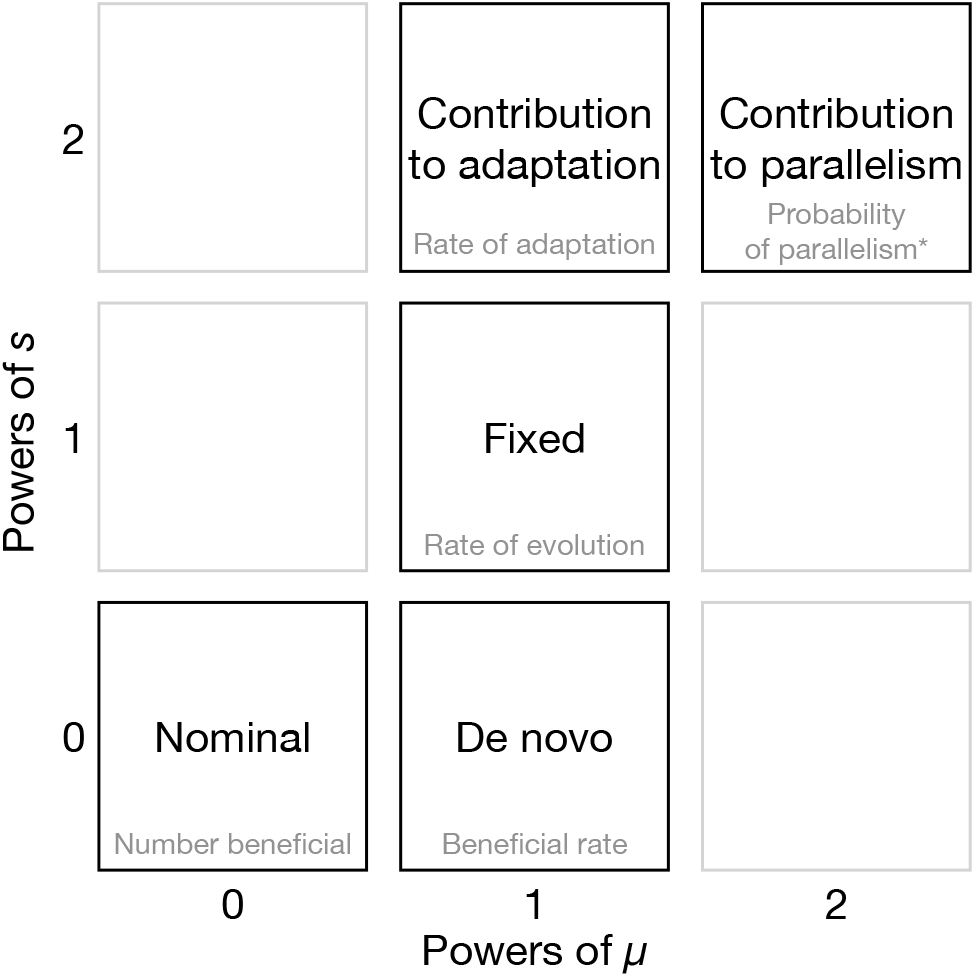
Further joint distributions obtained by size-biasing. Beyond the nominal, *de novo* and fixed distributions, are two additional joint distributions of evolutionary interest. Size-biasing the fixed distribution by the selection coefficients yields the distribution of contributions to the expected rate of adaptation. Size-biasing again by mutation rates results in a distribution where the probability for each beneficial mutation gives its relative contribution to the rate of parallel evolution. Values on the axes show how many times the nominal distribution is size-biased by *μ* or *s*. The gray text gives an evolutionary interpretation of the normalizing constant for each distribution. For the top right distribution, the normalization constant (*) is actually the rate of evolution squared, times the probability of parallelism (see the mathematical Appendix for details).

The graphical representation of these joint distributions shown in Figure 9 may be helpful for understanding some generic aspects of adaptation. For example, moving up the diagonal in Fig. 9, the fixed distribution is derived by doubly size-biasing the nominal, and the contribution to parallel adaptation is derived by doubly size-biasing the fixed distribution, so that the relationship between the contribution to parallelism and the fixed distribution is the same as the relationship between the fixed distribution and the nominal distribution.

Thus, just as the sign of the correlation between mutation rates and selection coefficients in the nominal determines whether the rate of evolution (the normalizing constant for the fixed distribution) is more or less than it would be if mutation rates and selection coefficients were uncorrelated in the nominal, the correlation in the fixed distribution determines whether the probability of parallelism (the normalizing constant for the contribution to parallelism) is increased or decreased relative to the case in which mutation rates and selection coefficients are uncorrelated among fixed mutations. As a consequence, if the fixed distribution shows a negative correlation (as we suggest is likely), this would tend to decrease the probability of parallelism.

A second general direction for further theoretical work is to explore the emergence of associations between mutation rates and selection coefficients under a broader range of conditions. Our treatment here is restricted to evolution via fixation of beneficial mutations in the SSWM regime where the probability of fixation is proportional to *s*, so that our treatment can be conducted by size-biasing according to *μ* and by *s*. More generally, however, evolution is not necessarily mutation-limited, and it includes deleterious, neutral, and beneficial changes, so that the chances of a given change are not guaranteed to be proportional to either *s* or *μ*. Under strict origin-fixation conditions (i.e. in the limit as *Nμ* → 0), and considering all classes of fitness effects—deleterious, neutral and beneficial—, the effect of mutation is proportional to *μ* and the effect of selection is given by the probability of fixation, which is a function of *s* and *N* (Kimura, 1962). As mutation supply increases, clonal interference comes into play and, where studied, this amplifies the effect of selection and diminishes the influence of mutation bias (e.g., Bailey et al. 2017; Cano et al. 2022; Gomez et al. 2020). Under the extreme condition that all possible variants are readily available in a large population, selection picks the winner and the chance of fixation is zero for all but the most fit variant (Bailey et al., 2017). Effects similar to what we observe here could occur under a variety of conditions, wherever effects of mutation and selection are both strong.

A final set of issues concerns what determines the association of mutation and fitness in the nominal, both in the proximate sense of the biological factors accounting for any association, and in the ultimate sense of the evolutionary dynamics that shape any mutation-fitness associations via these biological factors. Clearly this set of issues is more subtle and complex than is suggested by the textbook doctrine of random mutation (for analysis, see Merlin 2010; Razeto-Barry and Vecchi 2016; Stoltzfus 2021), which is often stated as a central tenet of the neo-Darwinian theory of evolution (Lenski and Mittler, 1993), though it only appears rarely as an explicit assumption in formal models (e.g., Chevin et al., 2010). First, it will be clear from our comments about sources of empirical data that actual measurements of the joint distribution of *s* and *μ* simply did not exist in any systematic form until very recently, so that there has been no empirical justification for any detailed quantitative claim about their relationship. Rather than being justified by systematic observations (or by biological principles), the randomness doctrine is perhaps best understood as a heuristic or guiding assumption for a neo-Darwinian research program (e.g., as argued in Stoltzfus 2021). Yet, if we choose to embrace the doctrine in this context, there are further issues. One issue is an ambiguity about whether randomness refers to the nominal, the *de novo*, or perhaps the underlying distribution from which the realized nominal distribution is drawn. Finally, the theory presented here raises the question of what, precisely, is the heuristic benefit of a randomness assumption. If there is no correlation in the nominal, this ensures that mutation biases do not increase the rate of evolution, but does not prevent an effect on the rate of adaptation, for reasons explained above. Likewise, if the lack of correlation refers to the *de novo*, this ensures that the average mutation rate among fixed mutations is equal to the average rate among *de novo* mutations, but does not prevent mutational biases from increasing either the rate of adaptation or the rate of evolution.

Meanwhile, diverse ideas about the evolution of mutation-fitness associations have been proposed, from relatively narrow and modest claims of adaptive amelioration lowering the mutation rate in functionally important regions (Martincoreña and Luscombe, 2013; Monroe et al., 2022), to the emergence of specialized mutation systems (e.g., cassette-shuffling systems) in the context of immune evasion or host-phage arms races (Foley 2015; Ch. 5 of Stoltzfus 2021), to ideas about “directed” or “smart” mutation systems (Roth et al., 2006). New thinking continues to emerge on these and related topics, e.g., Oman et al. (2022) explore the issue of what happens to the mutation rate and pattern as a genome evolves under context-dependent mutation. In adaptive walks, simply reversing mutation biases partway through the walk appears to improve adaptation by shifting the *de novo* distribution in favor of previously low rate beneficial mutations that have not yet had the chance to fix (Sane et al., 2022). For polygenic quantitative traits, continued evolution under correlated selection can lead to a shift in how the traits are encoded so that mutational variability aligns more closely with trait correlations favored by selection (Jones et al., 2014). Thus, a challenge for future theoretical work is to consider these ideas together in a common framework.

## Methods

### Empirical distributions for Dengue virus and TP53

We use data containing estimates of selection coefficients and mutation rates for all possible single point mutations in the Dengue genome, provided in Extended Table Data 1 (Dolan et al., 2021). The mutation rates provided in this table apply to the 12 types of context-independent single-nucleotide changes that are possible in a single stranded RNA genome such as the Dengue genome. In terms of the fitness measurements, we filtered the data to retain only beneficial variants for which the lower limit of the fitness confidence interval exceeds 1.0, using data for passage 7, replicate A, on human host cells. The final list contains 237 beneficial mutations.

With regard to TP53, Giacomelli et al. (2018) provide enrichment scores for the wild-type and all 7880 possible non-synonymous variants. To fully assess the potential for cancer-causing mutations, they combined results of multiple selection assays based on the effects of a second wild-type or null p53 allele (in otherwise isogenic human lung carcinoma cell), and the p53-activating agents nutlin3 and etoposide. This resulted in 3 assays designed to enrich for dominant-negative (wt with nutlin3), loss-of-function (null with nutlin3), or wt-like (null with etoposide) alleles. Following Giacomelli et al. (2018), we combined the estimated enrichments for the three assays using the formula *R*_all_ = *R*_(wt nutlin3)_ + *R*_(null nutlin3)_ - *R*_(null etoposide)_, and calculated selection coefficients B for all possible non-synonymous variants (including single- and multinucleotide variants) in terms of this combined enrichment score relative to the wild-type TP53 combined enrichment score: 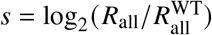.

We note that multiple studies have reported measures of cancer-causing potential for TP53 mutants based on deep mutational scanning (Giacomelli et al., 2018; Kotler et al., 2018; Staller et al., 2022). The results tend to be highly correlated (Carbonnier et al., 2020), but differ widely in scale. In the data from Giacomelli et al. (2018), most selection coefficients exceed 1, and the largest is 7, whereas other studies report a more modest range. This issue of scaling will not affect the shape of any correlation with mutation rates. However, in population-genetic simulations, the scale of selection coefficients matters for the chance of fixation and for the duration of fixation events, which influences clonal interference. Therefore, because the reported scale of selection coefficients may be artificially high, we use the reported selection coefficients as well as selection coefficients reduced 10-fold or 100-fold.

To estimate the mutation spectrum for TP53 we used a model based on data from the Pancancer Analysis of Whole Genomes database (Weinstein et al., 2013). Specifically, we queried a total of 28717344 whole genome single point somatic mutations to construct a trinucleotide context model (i.e., based on the identity of the nucleotides upstream and downstream of the focal nucleotide). For each possible mutation in each possible tri-nucleotide context we divided the observed frequency of that mutation within our dataset of somatic mutations by the genomic frequency of that tri-nucleotide context. The resulting vector with the rates for the 96 different mutation types is then normalized to yield fractional mutation rates that sum to 1 (Fig. S7). When a possible beneficial amino acid change can occur by multiple types of nucleotide mutations, we sum their rates.

For the clinically observed distribution of TP53 variants, we extracted the observed frequency of 639 different somatic mutations in TP53 in human tumors from the GENIE database of the American Association for Cancer Research Consortium AACR Project Genie Consortium et al. (2017). Note that this source aggregates data from different types of cancers, a heterogeneity that we ignore here.

We then integrated the observed frequencies for cancer mutations with the previously calculated selection coefficients and mutation rates. Note that we use different sources for (1) the selection coefficients (deep mutational scanning), (2) the mutation spectrum (Pancancer Analysis of Whole Genomes) and (3) the observed frequencies that we use to represent the fixed distribution (GENIE).

### Evolutionary simulations

We used SLiM v3.4 for the evolutionary simulations (Messer, 2013). We ran each simulation until a single sequence went to fixation (frequency > 0.95), recording any beneficial mutations in the fixed sequence. We repeated this process 1000 times per value of mutation supply *Nμ*_tot_ (*μ*_tot_ constant, *N* varies). Each of the simulations per replicate used the same initial haploid Wright-Fisher population, comprising *N* copies of the wild-type sequence of the Dengue virus genome or the TP53 coding region (with *N* ranging from 12 to 1.2 ×10^5^ or 100 to 10^6^, respectively). For Dengue virus, *μ*_tot_ is chosen to give the empirically estimated genome-wide beneficial mutation rate for the wild type genome *μ*_tot_ = 8.1 × 10^−6^. In the case of TP53 we use the empirical mutational signature model of relative rates for each mutation type, assuming *μ*_tot_ = 1.0 × 10^−6^. All sequences in the initial population were assigned a fitness of one. In each generation *t, N* sequences were chosen from the population at generation *t* - 1 with replacement and with a probability proportional to their fitness. The fitness effects assigned to each of the possible adaptive changes were taken from their respective datasets (single point mutations for Dengue and amino acid changes for TP53). Mutations continue to occur in derived alleles, so that secondary mutations are possible (with additive effects on fitness), although the sequence fixed in a replicate only very rarely has more than one mutation.

## Acknowledgments

We thank Dr. Patrick Dolan for help interpreting Dengue virus data. We thank Dr. Debra Kaiser and two reviewers for helpful comments on the manuscript. This work was made possible through the support of a grant from the John Templeton Foundation (grant #61782, D.M.M.) and from the Swiss National Science Foundation (grants #PP00P3_170604 and #310030_192541, J.L.P.). The opinions expressed in this publication are those of the authors and do not necessarily reflect the views of the John Templeton Foundation. D.M.M. also acknowledges additional support from an Alfred P. Sloan Research Fellowship and from the Simons Center for Quantitative Biology. The identification of any specific commercial products is for the purpose of specifying a protocol, and does not imply a recommendation or endorsement by the National Institute of Standards and Technology.

## Statement of Authorship

All authors were involved in conceptualization and design. All authors were involved in writing and editing the manuscript. BLG and DMM developed the mathematical theory of size-biasing and carried out associated simulations and visualizations. AVC designed and carried out population simulations and associated analysis and visualizations. AVC, AS and JLP gathered and assessed sources of data, including those from TP53 and Dengue virus.

## Data and code availability

Data and code used in this study (Gitschlag et al., 2023) are available in a GitHub repository at https://github.com/alejvcano/paradox (DOI: 10.5281/zenodo.7764132).

## Appendix Mathematical derivations

We begin by establishing some useful general results concerning size-biasing. Let *X* be a random variable taking non-negative values and. be another random variable such that *X* and *Y* can occur in *n* distinct pairs (*x*_1_, *y*_1_), …, (*x*_*n*_, *y*_*n*_). Furthermore let *X*^*^ and *Y*^*^ be obtained from the joint distribution of *X* and *Y* by size-biasing according to *X* so that the probability of (*x*_*i*_, *y*_*i*_) under the size-biased distribution is given by *x*_*i*_ *P* (*x*_*i*_, *y*_*i*_)/*E* (*X*). Then:

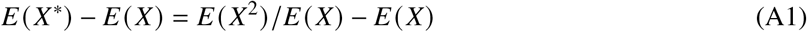

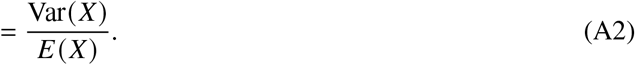

Similarly,

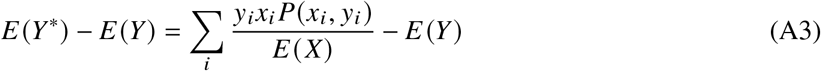

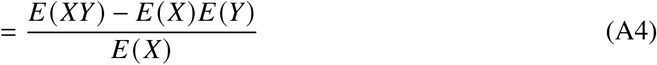

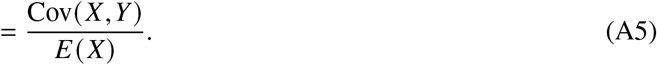

Equations 2 and 3 in the main text follow from these immediately by noting that the *de novo* distribution is obtained from the nominal distribution by size-biasing according to *μ*, and Equations 6 and 7 follow by noting that the fixed distribution is obtained from the *de novo* distribution by size-biasing with respect to *s*.

In fact, by repeatedly size-biasing with respect to *μ* and *s* we can develop a whole series of biologically relevant joint distributions of selection coefficients and mutation rates, where we size-bias *l* times with respect to *μ* and *m* times with respect to *s*. We begin by noting that often in evolution we are interested in quantities of the form 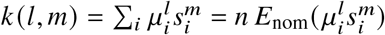. For example, *k* (0, 0) is the number of beneficial mutations and *k* (1, 0) is the total beneficial mutation rate. Assuming that the probability of fixation is a linear function of *s* (as explained in the main text), we also have that *k* (1, 1) is proportional to the rate of adaptive substitutions and *k* (1, 2) is proportional to the expected rate of fitness increase. Under this same approximation, we see that the probability of adaption via beneficial mutation *i* is given by *μ*_*i*_ *s*_*i*_/*k* (1, 1), so that the probability that two independent bouts of adaptation both proceed via fixation of *s* is given by (*μ*_*i*_ *s*_*i*_ /*k* (1, 1))^2^ and the total probability of parallel evolution (Chevin et al., 2010; Lenormand et al., 2016) is given by 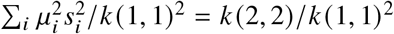. Thus we see that *k* (2, 2) is proportional to the probability of parallel evolution. Overall, size-biasing *l* times with respect to *μ* and *m* times with respect to *s* results in a probability distribution where mutation *i* occurs with probability 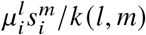, and this probability gives the proportional contribution of mutation *i* to *k* (*l, m*). For example, mutation *i* fixes at a rate proportional to *μ*_*i*_ *s*_*i*_ and increases fitness by *s*_*i*_ when it fixes, so that 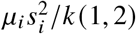 gives the relative contribution of mutation *i* to the total rate of adaptation.

Finally, we are interested in the sign of the correlation between *μ* and *s* in the distribution obtained by size-biasing *l* times with respect to *μ*and *m* times with respect to *s*. The correlation has the same sign as the covariance between *μ* and *s* in this distribution, where the covariance is given by:

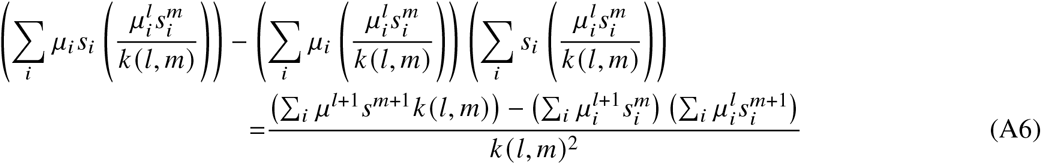

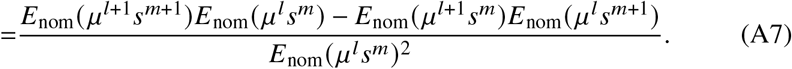

Noting that the denominator *E*_nom_(*μ* ^*l*^ *s*^*m*^)^2^ is always positive, we see that Cov_de novo_(*μ, s*) has the same sign as *E*_nom_(*μ* ^2^*s*)*E*_nom_(*μ* − *E*_nom_(*μ* ^2^)*E*_nom_(*μ s*) and Cov_fixed_(*μ, s*) has the same sign as *E*_nom_(*μ* ^2^*s* ^2^)*E*_nom_(*μ s*) − *E*_nom_(*μ* ^2^*s*)*E*_nom_(*μ s* ^2^) as stated in the main text.

### The special case of two selection coefficients and two mutation rates

In what follows we analyze the special case where there are only 2 possible selection coefficients and 2 possible mutation rates. Here, we alter the notation slightly as follows. Because the associations between mutation rates and selection coefficients depend on relative values, mutation rates are expressed as either 1 for the baseline or *B* where *B* > 1 for the elevated value, while selection coefficients are expressed as either 1 for the baseline value or *K* where *K* > 1 for the elevated value. Moreover, rather than indexing each mutation separately, we instead express the distributions in terms of the relative proportion of potential mutations with each unique pairwise combination of mutation rate and selection coefficient, so that, for example, 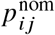 represents the probability that a mutation, sampled uniformly at random from a list of potential mutations, will have mutation rate *i* and selection coefficient *j*. We also write (for example) 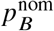 for the fraction of mutations in the nominal distribution with the larger mutation rate and 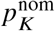 for the fraction of mutations in the nominal distribution with the larger selection coefficient.

Using this notation we can write Cov_nom_(*μ, s*) as:

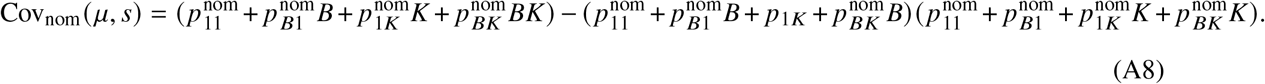

The above expression can be simplified by solving for one of the relative proportions in terms of the remaining three: 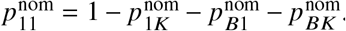. Substituting this value for 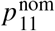 into the above expression, the covariance between mutation rate and selection coefficient is given by:

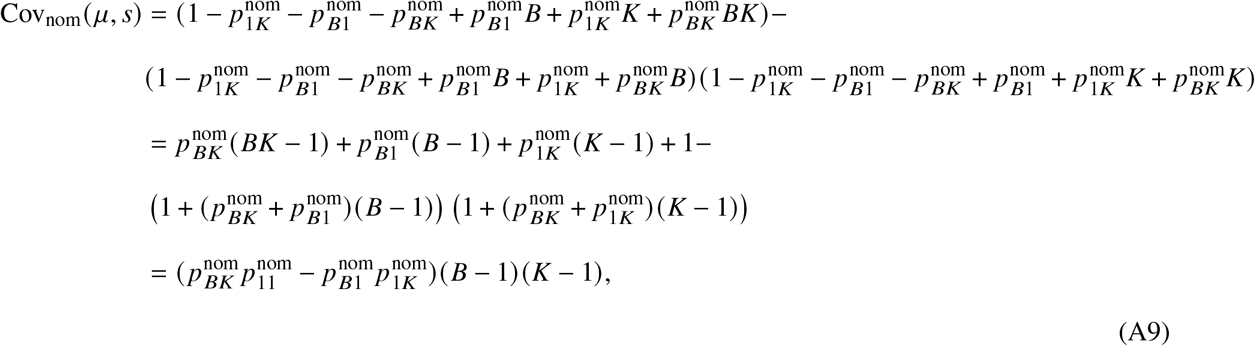

where the last line follows by expanding the product, applying the identity 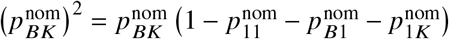, and then simplifying. The corresponding correlation coefficient is then given by dividing by the product of the standard deviations:

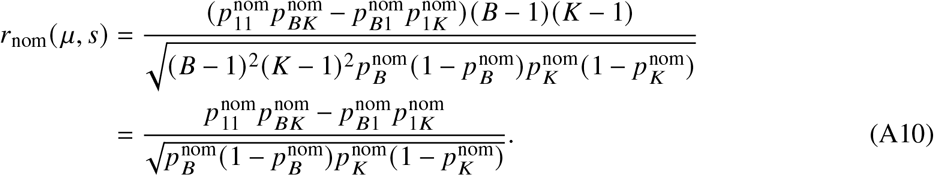

Finally, if we let 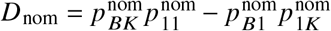 and note that *p*(1 - *p*) is the variance of a Bernoulli random variable with parameter *p*, then we obtain Equation 8 in the main text:

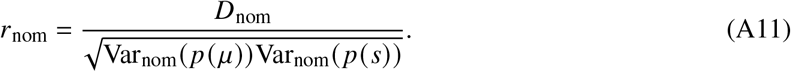

By making similar substitutions as those used for deriving the covariance among nominal mutations, we derive the following simplified expression for covariance of mutation rate and selection coefficient among *de novo* mutations when mutation rates and selection coefficients each take only two values:

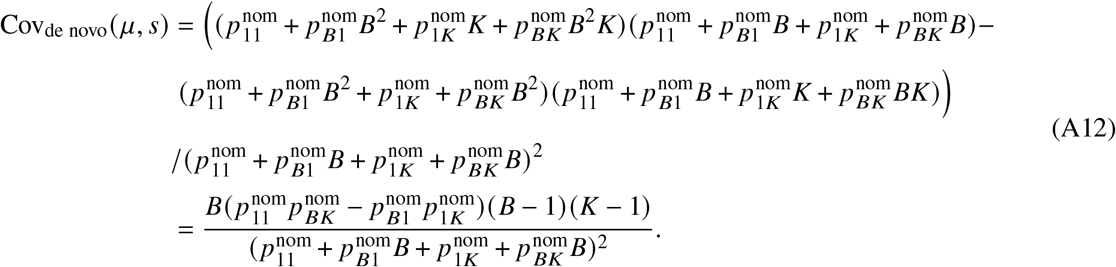

The correlation coefficient can be derived in similar fashion:

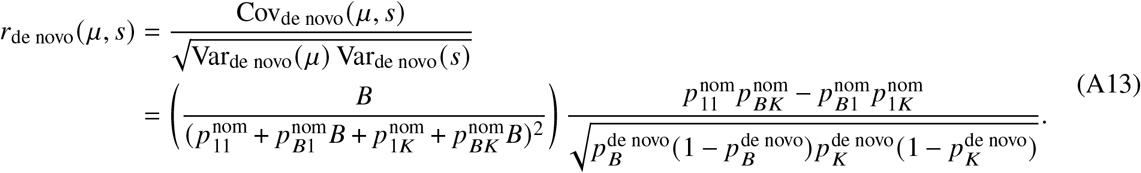

Finally, if we let 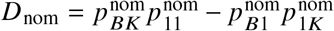 as before, then we obtain the following, simplified formula for the correlation coefficient among *de novo* mutations, corresponding to Equation 9 in the main text:

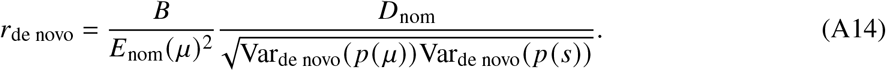

Making similar substitutions as those used for deriving each respective covariance for nominal and *de novo* mutations, we derive the following simplified covariance expression among fixed mutations:

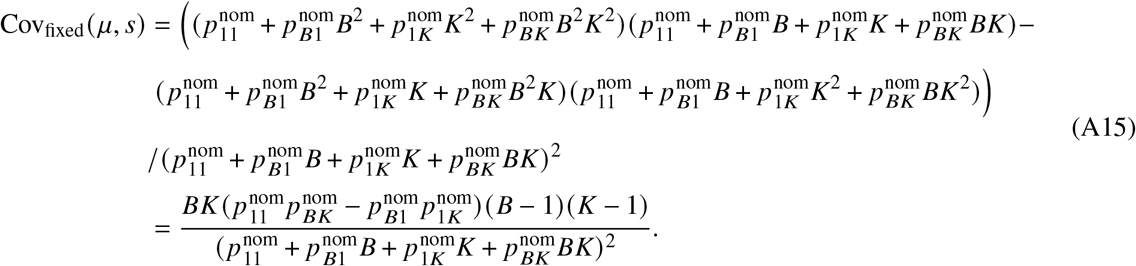

The correlation coefficient can be derived in similar fashion as before:

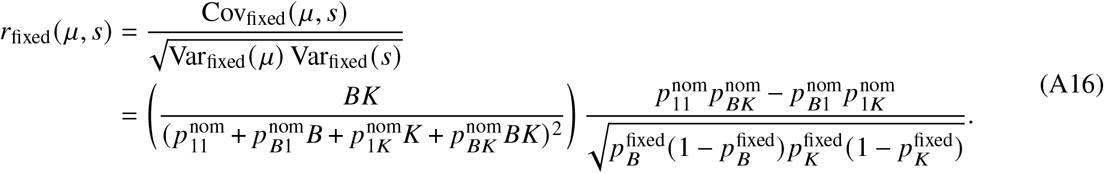

Finally, if we let 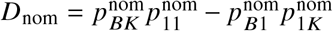 as before, then we obtain the following, simplified formula for the correlation coefficient among fixed mutations, corresponding to main text Equation 10:

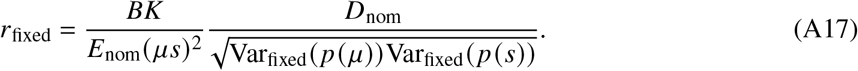

Although the above results are based on using the Pearson’s correlation as a measure of association, it is also interesting to ask about non-parametric measures of association such as Goodman-Kruskal’s *γ*(Goodman and Kruskal, 1954). For the special case with two distinct mutation rates and two distinct selection coefficients considered here, the probability of concordance for the nominal distribution is given by 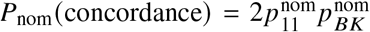 and the probability of discordance is given by 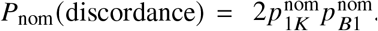. Thus, we have

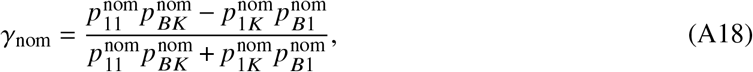

where we note that the denominator is positive and the numerator is simply *D*_nom_, so *r*_nom_ and *γ*_nom_ will have the same sign. Moving to the *de novo* distribution, and noting that the normalizing factors for the four probabilities all cancel out, we have

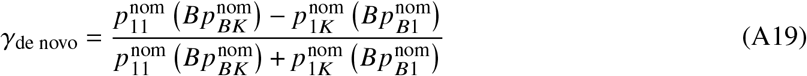

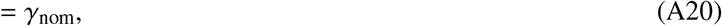

where the second equality holds because each product in the expression contains exactly one factor of *B* and so all these factors of *B* cancel out. Similarly, we have

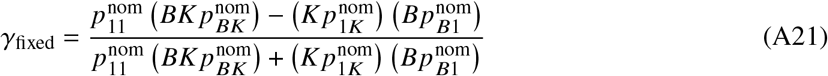

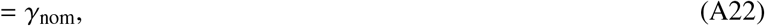

since each product contains exactly one factor of each of *B* and *K*. Thus for this special case we have *γ*_nom_ = *γ*_de novo_ = *γ*_fixed_ as stated in the main text.

## Literature Cited

AACR Project Genie Consortium, et al. 2017. Aacr project genie: powering precision medicine through an international consortium. Cancer discovery 7:818–831.

Acevedo, A., L. Brodsky, and R. Andino. 2014. Mutational and fitness landscapes of an rna virus revealed through population sequencing. Nature 505:686–690.

Arratia, R., L. Goldstein, and F. Kochman. 2019. Size bias for one and all. Probability Surveys 16:1–61.

Bailey, S. F., F. Blanquart, T. Bataillon, and R. Kassen. 2017. What drives parallel evolution?: How population size and mutational variation contribute to repeated evolution. Bioessays 39:1–9.

Cannataro, V. L., S. G. Gaffney, T. Sasaki, N. Issaeva, N. K. S. Grewal, J. R. Grandis, W. G. Yarbrough, B. Burtness, K. S. Anderson, and J. P. Townsend. 2019. Apobec-induced mutations and their cancer effect size in head and neck squamous cell carcinoma. Oncogene 38:3475–3487.

Cannataro, V. L., S. G. Gaffney, and J. P. Townsend. 2018. Effect sizes of somatic mutations in cancer. J Natl Cancer Inst 110:1171–1177.

Cano, A. V. B. Gitschlag, H. Rozhonová, A. Stoltzfus, D. M. McCandlish, and J. L. Payne. 2023. Mutation bias and the predictability of evolution. Phil. Trans. R. Soc. B 378:20220055.

Cano, A. V., H. Rozhonová, A. Stoltzfus, D. M. McCandlish, and J. L. Payne. 2022. Mutation bias shapes the spectrum of adaptive substitutions. Proc Natl Acad Sci U S A 119.

Carbonnier, V., B. Leroy, S. Rosenberg, and T. Soussi. 2020. Comprehensive assessment of tp53 loss of function using multiple combinatorial mutagenesis libraries. Sci Rep 10:20368.

Chevin, L. M., G. Martin, and T. Lenormand. 2010. Fisher’s model and the genomics of adaptation: restricted pleiotropy, heterogenous mutation, and parallel evolution. Evolution 64:3213–31.

Couce, A., A. Rodríguez-Rojas, and J. Blazquez. 2015. Bypass of genetic constraints during mutator evolution to antibiotic resistance. Proceedings of the Royal Society London B 282:20142698.

Darwin, C. 1868. Variation of Animals and Plants under Domestication, vol. II.

Murray London. Diss G., and B. Lehner. 2018. The genetic landscape of a physical interaction. Elife 7.

Dolan, P. T., S. Taguwa, M. A. Rangel, A. Acevedo, T. Hagai, R. Andino, and J. Frydman. 2021. Principles of dengue virus evolvability derived from genotype-fitness maps in human and mosquito cells. eLife 10:e61921.

Eyre-Walker, A., and P. D. Keightley. 2007. The distribution of fitness effects of new mutations. Nat Rev Genet 8:610–8.

Foley, J. 2015. Mini-review: Strategies for variation and evolution of bacterial antigens. Comput Struct Biotechnol J 13:407–16.

Gerrish, P. J., and R. E. Lenski. 1998. The fate of competing beneficial mutations in an asexual population. Genetica 102:127–144.

Giacomelli, A. O., X. Yang, R. E. Lintner, J. M. McFarland, M. Duby, J. Kim, T. P. Howard, D. Y. Takeda, S. H. Ly, E. Kim, et al. 2018. Mutational processes shape the landscape of tp53 mutations in human cancer. Nature genetics 50:1381–1387.

Gillespie, J. H. 1994. The Causes of Molecular Evolution. Oxford University Press.

Gitschlag, B., A. V. Cano, J. L. Payne, D. M. McCandlish, and A. Stoltzfus. 2023. Data from: Mutation and selection induce correlations between selection coefficients and mutation rates.

Gomez, K., J. Bertram, and J. Masel. 2020. Mutation bias can shape adaptation in large asexual populations experiencing clonal interference. Proceedings of the Royal Society B: Biological Sciences 287:20201503.

Good, B. H., I. M. Rouzine, D. J. Balick, O. Hallatschek, and M. M. Desai. 2012. Distribution of fixed beneficial mutations and the rate of adaptation in asexual populations. Proc Natl Acad Sci U S A 109:4950–5.

Goodman, L. A., and W. H. Kruskal. 1954. Measures of association for cross classifications. Journal of the American Statistical Association 49:732–764.

Hainaut, P., and G. P. Pfeifer. 2016. Somatic tp53 mutations in the era of genome sequencing. Cold Spring Harbor Perspectives in Medicine 6:a026179.

Haldane, J. 1927. A mathematical theory of natural and artificial selection. v. selection and mutation. Proc. Cam. Phil. Soc. 26:220–230.

Hodgkinson, A., and A. Eyre-Walker. 2011. Variation in the mutation rate across mammalian genomes. Nat Rev Genet 12:756–766.

Jones, A. G., R. Burger, and S. J. Arnold. 2014. Epistasis and natural selection shape the mutational architecture of complex traits. Nat Commun 5:3709.

Kassen, R., and T. Bataillon. 2006. Distribution of fitness effects among beneficial mutations before selection in experimental populations of bacteria. Nat Genet 38:484–8.

Katju, V., and U. Bergthorsson. 2019. Old trade, new tricks: Insights into the spontaneous mutation process from the partnering of classical mutation accumulation experiments with high-throughput genomic approaches. Genome Biol Evol 11:136–165.

Kimura, M. 1962. On the probability of fixation of mutant genes in a population. Genetics 47:713–9.

King, J., and T. Jukes. 1969. Non-darwinian evolution. Science 164:788–797.

Kotler, E., O. Shani, G. Goldfeld, M. Lotan-Pompan, O. Tarcic, A. Gershoni, T. A. Hopf, D. S. Marks, M. Oren, and E. Segal. 2018. A systematic p53 mutation library links differential functional impact to cancer mutation pattern and evolutionary conservation. Mol Cell 71:178–190 e8.

Leighow, S., C. Liu, H. Inam, B. Zhao, and J. Pritchard. 2020. Multi-scale predictions of drug resistance epidemiology identify design principles for rational drug design. Cell Reports 30:3951–3963.

Lenormand, T., L. M. Chevin, and T. Bataillon. 2016. Parallel evolution: what does it (not) tell us and why is it (still) interesting? Pages 196–220 in G. Ramsey and C. Pence, eds. Chance in Evolution. University of Chicago Press, Chicago.

Lenski, R. E., and J. E. Mittler. 1993. The directed mutation controversy and neo-darwinism. Science 259:188–94.

Láruson, A. J., S. Yeaman, and K. E. Lotterhos. 2020. The importance of genetic redundancy in evolution. Trends Ecol Evol 35:809–822.

Macdonald, C. B., D. Nedrud, P. R. Grimes, D. Trinidad, J. S. Fraser, and W. Coyote-Maestas. 2022. Deep insertion, deletion, and missense mutation libraries for exploring protein variation in evolution, disease, and biology. bioRxiv page 2022.07.26.501589.

Maclean, C., G. Perron, and A. Gardner. 2010. Diminishing returns from beneficial mutations and pervasive epistasis shape the fitness landscape for Rifampicin resistance in Pseudomonas aeruginosa. Genetics 186:1345–54.

Martincoreña, I., and N. M. Luscombe. 2013. Non-random mutation: the evolution of targeted hypermutation and hypomutation. BioEssays : news and reviews in molecular, cellular and developmental biology 35:123–30.

McCandlish, D., and A. Stoltzfus. 2014. Modeling evolution using the probability of fixation: history and implications. Quarterly Review of Biology 89:225–252.

Merlin, F. 2010. Evolutionary Chance Mutation: A Defense of the Modern Synthesis’ Consensus View. Philosophy and Theory in Biology 2.

Messer, P. W. 2013. SLiM: Simulating evolution with selection and linkage. Genetics.

Monroe, J. G., T. Srikant, P. Carbonell-Bejerano, C. Becker, M. Lensink, M. Exposito-Alonso, M. Klein, J. Hildebrandt, M. Neumann, D. Kliebenstein, M. L. Weng, E. Imbert, J. Ågren, M. T. Rutter, C. B. Fenster, and D. Weigel. 2022. Mutation bias reflects natural selection in arabidopsis thaliana. Nature 602:101–105.

Neher, R. A. 2013. Genetic draft, selective interference, and population genetics of rapid adaptation. Annual review of Ecology, evolution, and Systematics 44:195–215.

Oman, M., A. Alam, and R. W. Ness. 2022. How sequence context-dependent mutability drives mutation rate variation in the genome. Genome Biol Evol 14.

Orr, H. A. 2003. The distribution of fitness effects among beneficial mutations. Genetics 163:1519–26.

Patwa, Z., and L. M. Wahl. 2008. The fixation probability of beneficial mutations. J R Soc Interface 5:1279–89.

Queller, D. C. 2017. Fundamental theorems of evolution. The American Naturalist 189:345–353.

Razeto-Barry, P., and D. Vecchi. 2016. Mutational randomness as conditional independence and the experimental vindication of mutational Lamarckism. Biol Rev Camb Philos Soc.

Rokyta, D., P. Joyce, S. Caudle, and H. Wichman. 2005. An empirical test of the mutational landscape model of adaptation using a single-stranded DNA virus. Nature Genetics 37:441–444.

Ross, S. M. 2014. Introduction to probability models. Academic press.

Roth, J. R., E. Kugelberg, A. B. Reams, E. Kofoid, and D. I. Andersson. 2006. Origin of mutations under selection: the adaptive mutation controversy. Annu Rev Microbiol 60:477–501.

Ruelens, P., and J. de Visser. 2021. Clonal interference and mutation bias in small bacterial populations in droplets. Genes (Basel) 12.

Sackman, A. M., L. W. McGee, A. J. Morrison, J. Pierce, J. Anisman, H. Hamilton, S. Sanderbeck, C. Newman, and D. R. Rokyta. 2017. Mutation-driven parallel evolution during viral adaptation. Molecular Biology and Evolution 34:3243–3253.

Sane, M., G. D. Diwan, B. A. Bhat, L. M. Wahl, and D. Agashe. 2022. Shifts in mutation spectra enhance access to beneficial mutations. bioRxiv page 2020.09.05.284158.

Sanjuan, R., A. Moya, and S. F. Elena. 2004. The distribution of fitness effects caused by single-nucleotide substitutions in an rna virus. Proc Natl Acad Sci U S A 101:8396–401.

Schenk, M. F., M. P. Zwart, S. Hwang, P. Ruelens, E. Severing, J. Krug, and J. de Visser. 2022. Population size mediates the contribution of high-rate and large-benefit mutations to parallel evolution. Nat Ecol Evol 6:439–447.

Schrider, D. R., J. N. Hourmozdi, and M. W. Hahn. 2011. Pervasive multinucleotide mutational events in eukaryotes. Curr Biol 21:1051–4.

Smith, N. G., M. T. Webster, and H. Ellegren. 2003. A low rate of simultaneous double-nucleotide mutations in primates. Mol Biol Evol 20:47–53.

Staller, M. V., E. Ramirez, S. R. Kotha, A. S. Holehouse, R. V. Pappu, and B. A. Cohen. 2022. Directed mutational scanning reveals a balance between acidic and hydrophobic residues in strong human activation domains. Cell Syst 13:334–345 e5.

Stoltzfus, A. 2021. Mutation, Randomness and Evolution. Oxford.

Stoltzfus, A., and D. M. McCandlish. 2017. Mutational biases influence parallel adaptation. Molecular Biology and Evolution 34:2163–2172.

Stoltzfus, A., and R. W. Norris. 2016. On the causes of evolutionary transition:transversion bias. Mol Biol Evol 33:595–602.

Storz, J. F., C. Natarajan, A. V. Signore, C. C. Witt, D. M. McCandlish, and A. Stoltzfus. 2019. The role of mutation bias in adaptive molecular evolution: insights from convergent changes in protein function. Philosophical Transactions of the Royal Society B: Biological Sciences 374:20180238.

Thyagarajan, B., and J. D. Bloom. 2014. The inherent mutational tolerance and antigenic evolvability of influenza hemagglutinin. Elife page e03300.

Watson, C. J., A. Papula, G. Y. Poon, W. H. Wong, A. L. Young, T. E. Druley, D. S. Fisher, and J. R. Blundell. 2020. The evolutionary dynamics and fitness landscape of clonal hematopoiesis. Science 367:1449–1454.

Weinstein, J. N., E. A. Collisson, G. B. Mills, K. R. Shaw, B. A. Ozenberger, K. Ellrott, I. Shmulevich, C. Sander, and J. M. Stuart. 2013. The cancer genome atlas pan-cancer analysis project. Nature genetics 45:1113–1120.

Wu, N. C., A. P. Young, L. Q. Al-Mawsawi, C. A. Olson, J. Feng, H. Qi, S. H. Chen, I. H. Lu, C. Y. Lin, R. G. Chin, H. H. Luan, N. Nguyen, S. F. Nelson, X. Li, T. T. Wu, and R. Sun. 2014. High-throughput profiling of influenza a virus hemagglutinin gene at single-nucleotide resolution. Sci Rep 4:4942.

