## Supplementary materials for "Mutation and selection induce correlations between selection coefficients and mutation rates"

Bryan L. Gitschlag[1,†]

Alejandro V. Cano[2,3,†]

Joshua L. Payne[2,3]

David M. McCandlish[1,*]

Arlin Stoltzfus[4,5,*]

1. Simons Center for Quantitative Biology, Cold Spring Harbor Laboratory, Cold Spring Harbor, NY, USA;
2. Institute of Integrative Biology, ETH, Zurich, Switzerland;
3. Swiss Institute of Bioinformatics, Lausanne, Switzerland;
4. Office of Data and Informatics, Material Measurement Laboratory, NIST, Gaithersburg, MD;
5. Institute for Bioscience and Biotechnology Research, Rockville, USA.

† These authors contributed equally

to appear in *The American Naturalist*

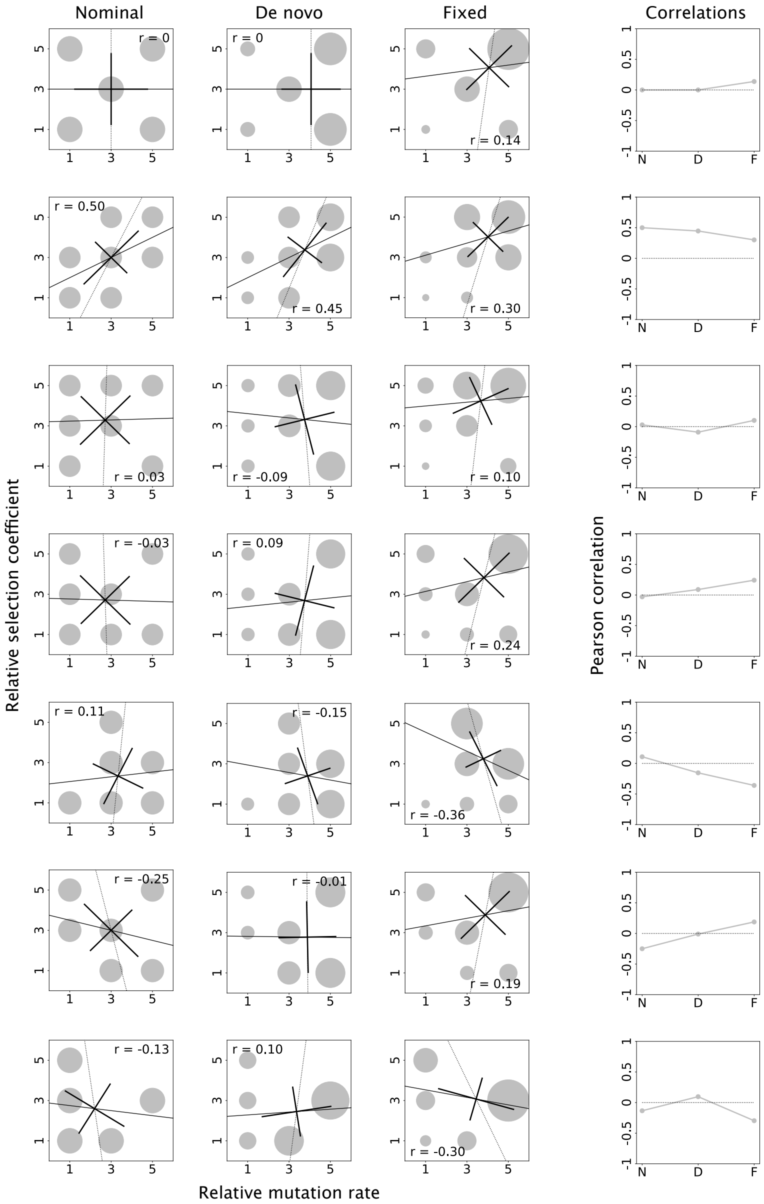


**Fig. S1. Correlations of mutation rates and selection coefficients under some alternative models to Fig. 2** As for Fig. 2, the columns show the nominal, *de novo* and fixed distributions, followed by a plot of Pearson's $r$ for the three distributions (designated by N, D and F). Here $\mu$ and $s$ each take on only 3 possible values. Values were chosen to illustrate (together with the bottom 2 rows of Fig. 2) that $r$ can take any pattern of signs across the nominal, *de novo*, and fixed. Regression lines and principal component axes as in Figure 2.

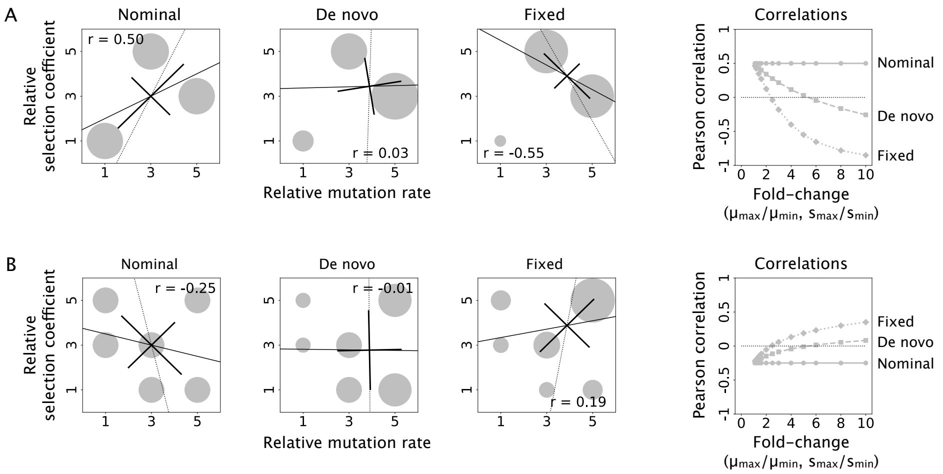


**Fig. S2. Effect of the magnitude of size-biasing. A.** Correlations between mutation rate and selection coefficient among nominal, *de novo*, and fixed distributions in a case where we see a sign-change in the correlation from positive in the nominal distribution to negative in the fixed distribution. Left to right: the nominal, *de novo*, and fixed distributions when mutation rate and selection coefficient each vary by 5-fold between their highest and lowest respective values (same as Fig. 2, fourth row), followed by the correlations between mutation rate and selection coefficient for the same 3-mutation-class distribution shown to the left, plotted as a function of the range, or fold-change, between highest and lowest value of mutation rate and selection coefficient ($\mu_{\max} = s_{\max}$ and $\mu_{\min} = s_{\min}$ throughout but $\mu_{\max}/\mu_{\min} = s_{\max}/s_{\min}$ is allowed to vary). Regression lines and principal component axes as in Figure 2. **B.** Correlations between mutation rate and selection coefficient among nominal, *de novo*, and fixed mutational distributions in a case where we see a sign-change in the correlation from negative in the nominal distribution to positive in the fixed distribution. Left to right: similar to panel **A**, the nominal, *de novo*, and fixed distributions when mutation rate and selection coefficient each vary by 5-fold between their highest and lowest respective values (same as Fig. S1, second row), followed by the correlations between mutation rates and selection coefficients for the same 3-mutation-class distribution shown to the left, plotted as a function of the range, or fold-change, between highest and lowest value of mutation rate and selection coefficient.

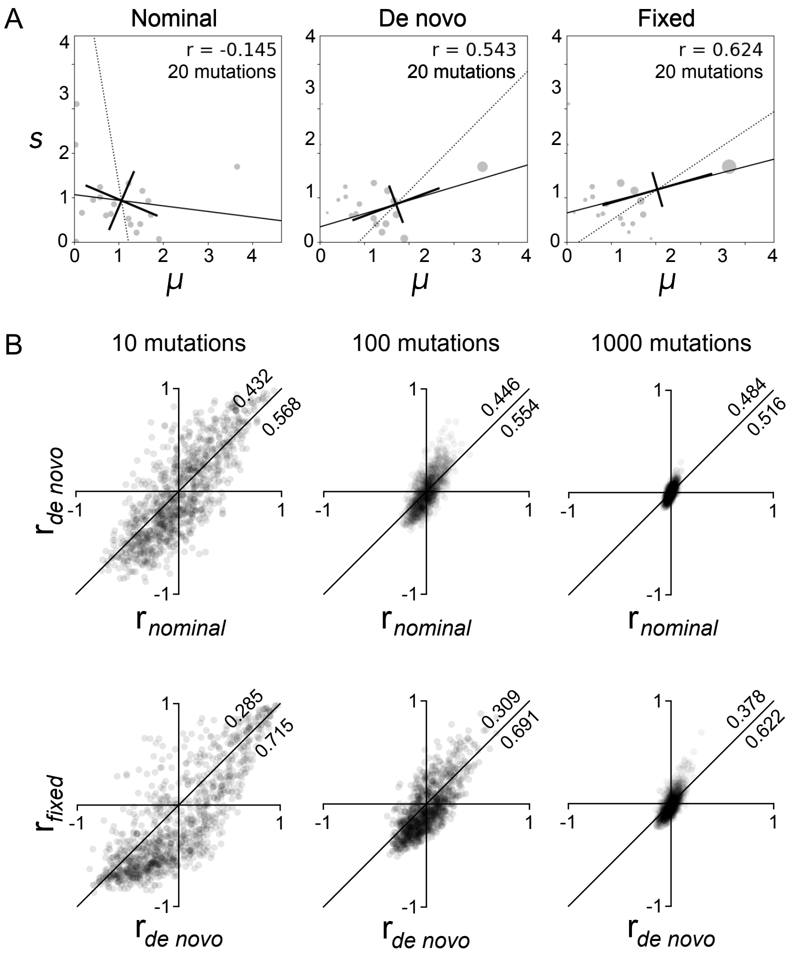


**Fig. S3. Effects of finite mutational target size and alternative nominal distributions on changes between joint distributions with respect to the correlation between mutation rates and selection coefficients in *de novo* and fixed distributions. A.** Similar to Fig. 3A, but using an example of a randomly generated nominal distribution representing the *less common case* in which the presence of a possible beneficial mutation with both high $\mu$ and high $s$ induces a strong positive correlation in the fixed distribution. **B.** Similar to Fig. 3B but showing the bivariate scatterplots of $r$ (Pearson's correlation) for *de novo* vs nominal distributions (top row) and fixed vs *de novo* distributions (bottom row), each for 1000 samples of 10, 100, and 1000 beneficial mutations.

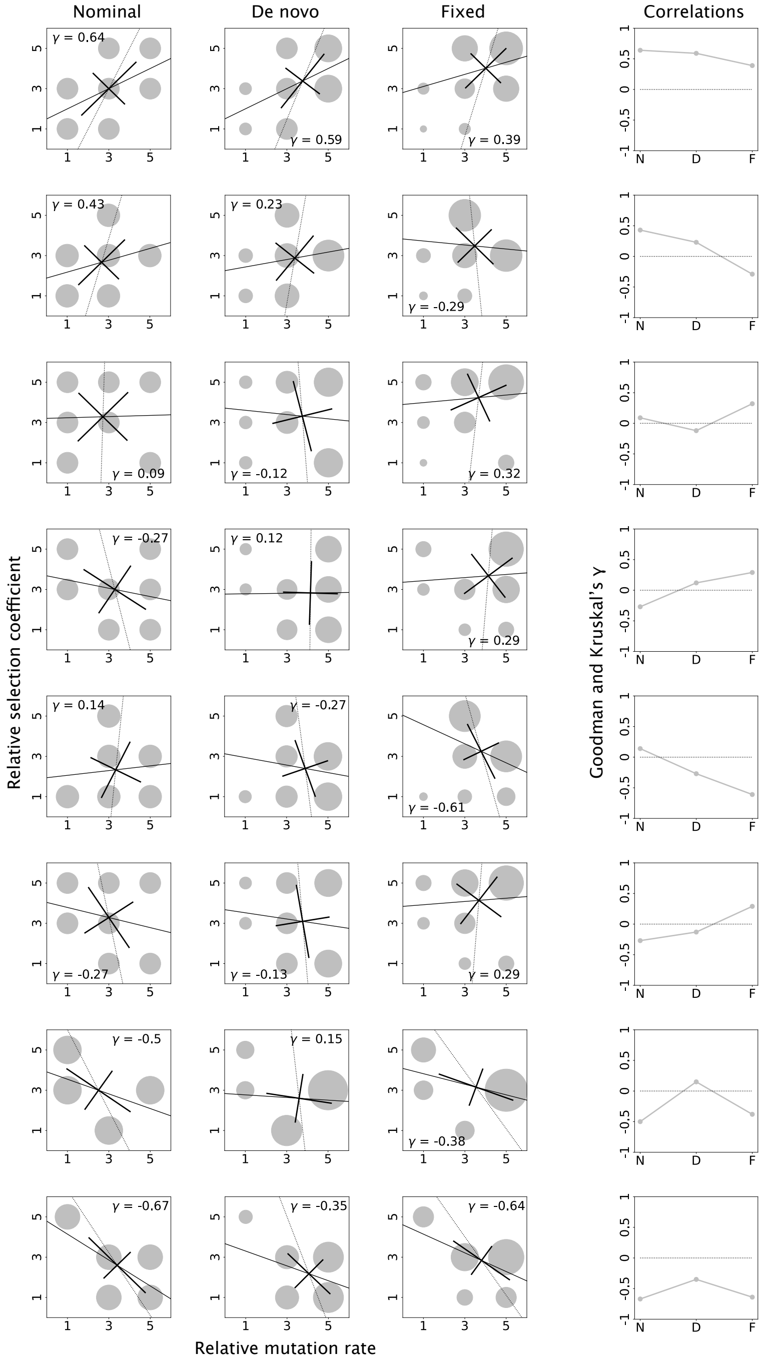


**Fig. S4. Non-parametric measures of association between mutation rate and selection coefficient.** Various choices of the nominal distribution illustrate that, as for the Pearson's correlation $r$ shown in Fig. S1, Goodman-Kruskal's $\gamma$ can take any pattern of signs across the nominal, *de novo*, and fixed distributions. Note that the sign of $r$ and the sign of the slope of the regression line need not agree with the sign of $\gamma$. Regression lines and principal component axes as in Figure 2.

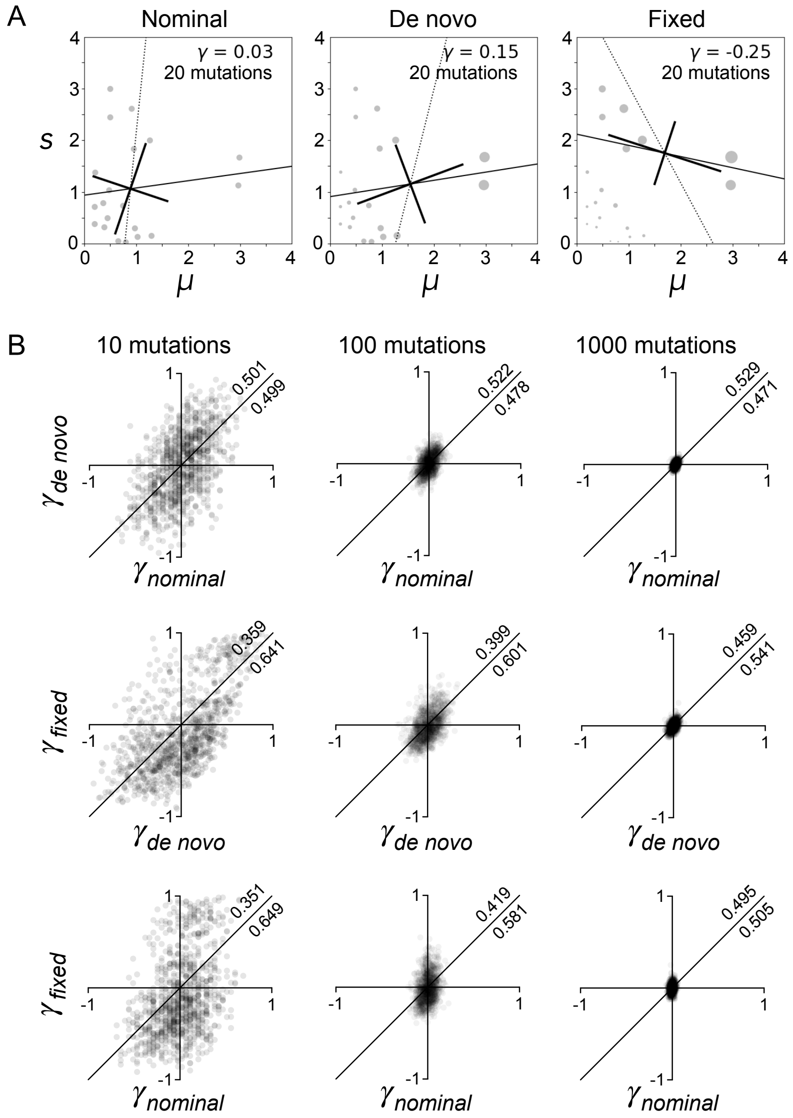


**Fig. S5. Finite mutational target sizes can change the direction of the non-parametric association between mutation rates and selection coefficients.** Representative nominal, *de novo*, and fixed distributions generated by sampling 20 mutations (top row) or 100 mutations (bottom row) from a joint distribution based on two independent exponential probability distributions (similar to Fig S3), to illustrate that Goodman-Kruskal's $\gamma$ can undergo sign-changes across the nominal, *de novo*, and fixed distributions. Notice that, similar to Pearson's $r$, Goodman-Kruskal's $\gamma$ tends to be more negative in the fixed distribution than it is in the nominal distribution.

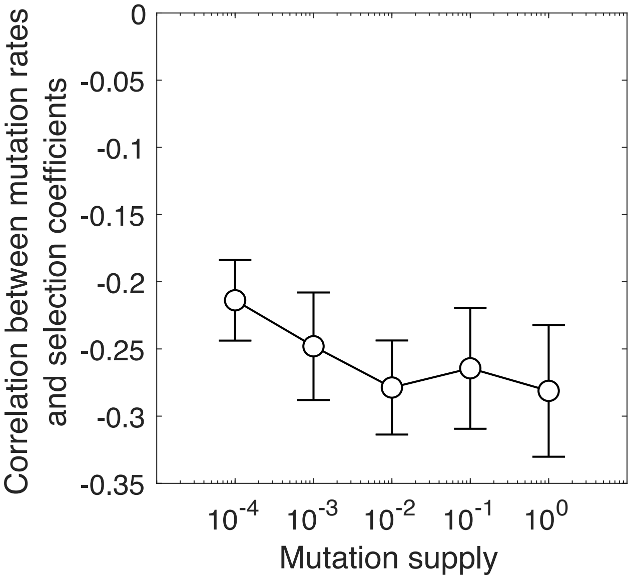


**Fig. S6. Goodman-Kruskal's $\gamma$ statistic for simulated fixed distributions based on the Dengue dataset.** Goodman-Kruskal's $\gamma$ statistic between mutation rates and selection coefficients for the distribution of fixed mutations in the Dengue dataset, as a function of mutation supply ($N\mu_{\text{tot}}$). Error bars depict 95% bootstrap confidence intervals based on $10^3$ bootstrap samples.

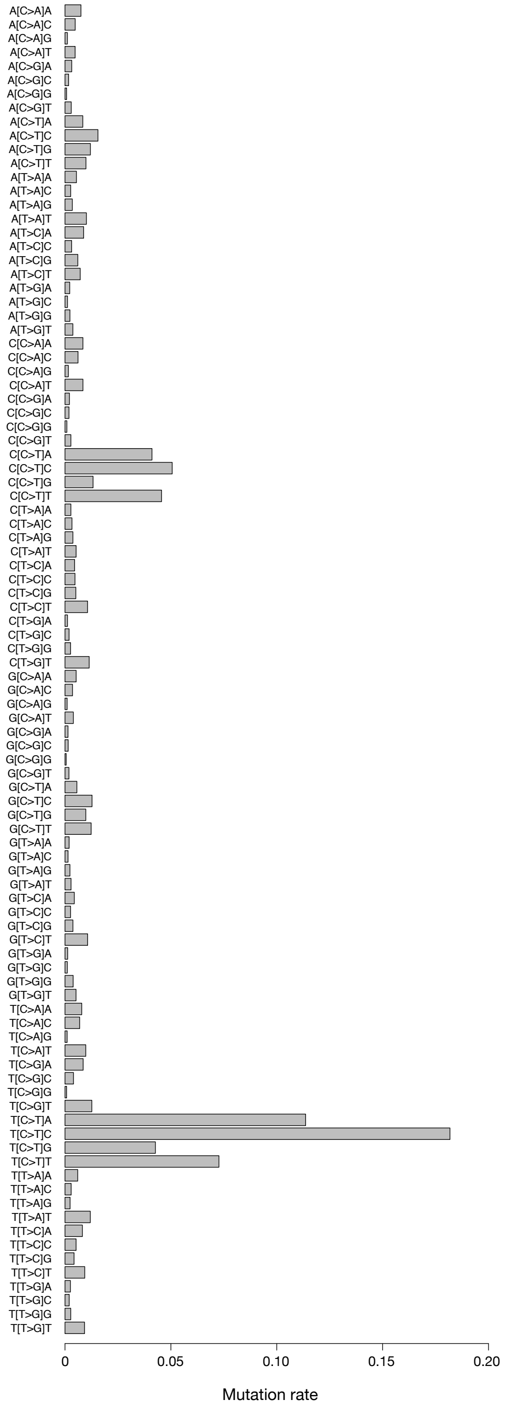


**Fig. S7. Context model for TP53 mutation rates.** We used $28,717,344$ single point somatic mutations taken from whole genome sequences in the Pancancer Analysis of Whole Genomes database (Weinstein et al., 2013) to estimate a mutational spectrum that incorporates the trinucleotide context, namely, the mutation rates specified by the identity of the mutated nucleotide and the immediately flanking nucleotides.

| $\mu_{tot}$ | PopSize | $\bar{s}$ | $r$ | 95% CI |
|---|---|---|---|---|
| 1.00E-06 | 1.00E+02 | 1.7 | 0.11 | [0.095,0.124] |
| 1.00E-06 | 1.00E+03 | 1.7 | 0.25 | [0.237,0.262] |
| 1.00E-06 | 1.00E+04 | 1.7 | 0.38 | [0.373,0.390] |
| 1.00E-05 | 1.00E+02 | 1.7 | 0.22 | [0.209,0.232] |
| 1.00E-05 | 1.00E+03 | 1.7 | 0.43 | [0.424,0.437] |
| 1.00E-05 | 1.00E+04 | 1.7 | 0.35 | [0.339,0.362] |
| 1.00E-04 | 1.00E+02 | 1.7 | 0.39 | [0.386,0.395] |
| 1.00E-04 | 1.00E+03 | 1.7 | 0.37 | [0.362,0.380] |
| 1.00E-04 | 1.00E+04 | 1.7 | 0.27 | [0.258,0.281] |
| 1.00E-06 | 1.00E+02 | 0.17 | 0.15 | [0.136,0.164] |
| 1.00E-06 | 1.00E+03 | 0.17 | 0.28 | [0.267,0.293] |
| 1.00E-06 | 1.00E+04 | 0.17 | 0.41 | [0.405,0.417] |
| 1.00E-05 | 1.00E+02 | 0.17 | 0.27 | [0.259,0.282] |
| 1.00E-05 | 1.00E+03 | 0.17 | 0.45 | [0.445,0.456] |
| 1.00E-05 | 1.00E+04 | 0.17 | 0.38 | [0.372,0.390] |
| 1.00E-04 | 1.00E+02 | 0.17 | 0.42 | [0.414,0.427] |
| 1.00E-04 | 1.00E+03 | 0.17 | 0.44 | [0.435,0.445] |
| 1.00E-04 | 1.00E+04 | 0.17 | 0.33 | [0.322,0.340] |
| 1.00E-06 | 1.00E+02 | 0.017 | 0.07 | [-0.050,0.091] |
| 1.00E-06 | 1.00E+03 | 0.017 | 0.19 | [0.176,0.205] |
| 1.00E-06 | 1.00E+04 | 0.017 | 0.27 | [0.258,0.282] |
| 1.00E-05 | 1.00E+02 | 0.017 | 0.16 | [0.142,0.177] |
| 1.00E-05 | 1.00E+03 | 0.017 | 0.29 | [0.277,0.302] |
| 1.00E-05 | 1.00E+04 | 0.017 | 0.31 | [0.301,0.320] |
| 1.00E-04 | 1.00E+02 | 0.017 | 0.26 | [0.249,0.270] |
| 1.00E-04 | 1.00E+03 | 0.017 | 0.28 | [0.265,0.294] |
| 1.00E-04 | 1.00E+04 | 0.017 | 0.35 | [0.338,0.361] |

**Table S1. The correlation between the observed and simulated versions of the fixed distribution depends on $N$, $\mu_{tot}$ and the scaling of relative selection coefficients**. Shown are the correlation coefficients $r$ with 95% bootstrap confidence intervals based on $10^6$ bootstrap samples.
